# Attenuated transcriptional response to pro-inflammatory cytokines in schizophrenia hiPSC-derived neural progenitor cells

**DOI:** 10.1101/2021.12.15.472107

**Authors:** Anjali Bhat, Haritz Irizar, Pooja Raval, Rodrigo R.R. Duarte, Lucia Dutan Polit, Timothy Powell, P.J. Michael Deans, Carol Shum, Roland Nagy, Grainne McAlonan, Conrad O. Iyegbe, Jack Price, Elvira Bramon, Sagnik Bhattacharyya, Anthony C. Vernon, Deepak P. Srivastava

## Abstract

Maternal immune activation (MIA) during prenatal development is an environmental risk factor for psychiatric disorders including schizophrenia (SZ). This link is particularly well established in the case of schizophrenia. Converging lines of evidence from human and animal model studies have suggested that elevated cytokine levels in the maternal and fetal compartments are an important indication of the mechanisms driving this association. However, there is variability in susceptibility to the psychiatric risk conferred by MIA, likely influenced by genetic factors. How MIA interacts with a genetic profile susceptible to SZ is challenging to test in animal models. To address this gap, we examined whether differential gene expression responses occur in forebrain-lineage neural progenitor cells (NPCs) derived from human induced pluripotent stem cells (hiPSC) generated from three individuals with a diagnosis of schizophrenia and three healthy controls. Following acute (24 hour) treatment with either interferon-gamma (IFNγ; 25 ng/μl) or interleukin (IL)-1β (10 ng/μl), we identified, by RNA sequencing, 3380 differentially expressed genes (DEGs) in the IFNγ-treated control lines (compared to untreated controls), and 1980 DEGs in IFNγ-treated SZ lines (compared to untreated SZ lines). Of these, 359 genes responded differently to IFNγ treatment in SZ relative to control lines, which were enriched for genes differentially expressed in adult SZ cases and in pathways related to the regulation of antigen processing, neuronal post-synaptic density, and the presynapse. There were no differentially expressed genes in the IL-1β-treatment conditions after Benjamini-Hochberg correction. Gene set enrichment analysis however showed that IL-1β impacts immune function and neuronal differentiation. Collectively, our data suggest that a) SZ NPCs show an attenuated transcriptional response to IFNγ treatment compared to control NPCs; b) IL-1β may be less relevant to NPC cultures than IFNγ and c) the genes differentially regulated in schizophrenia lines are primarily mitochondrial, loss-of-function, pre- and post-synaptic genes. Our findings particularly highlight early synaptic development as a key target for future research into the mechanisms of association between maternal immune activation and schizophrenia risk.

**Highlights:**

• There is a significant transcriptional response to IFNγ treatment in human induced cortical neural progenitor cells (NPCs)
• Differential expression implicates mitochondrial complex genes in this IFNγ-dependent response, which are underexpressed in response to treatment in schizophrenia (SZ) NPCs.
• SZ NPCs do not upregulate pre- and post-synaptic genes in response to IFNγ as much as control NPCs
• IL-1β may be less relevant to NPC cultures than IFNγ due to low IL1 receptor expression in NPCs.

## Introduction

Activation of the maternal immune response during pregnancy is a known risk factor for neurodevelopmental disorders – especially autism and schizophrenia – in the offspring [1–5]. Although the precise molecular mechanisms driving this association remain unclear, exposure of the developing fetal brain to pro-inflammatory cytokines remains a promising candidate for study [2, 6–8]. Cytokines are cell signalling proteins that help immune cells to form coordinated responses to infection. Whilst their function in the peripheral immune system is well documented, there is growing evidence that cytokines also play an important role in brain development and that maternally-derived cytokines could affect the developing foetal brain [2, 6–8]. Emerging evidence from human studies suggest that elevated levels of canonical pro-inflammatory cytokines such as interferon gamma (IFNγ) [2, 9] and interleukin-1 beta (IL-1β) [7, 10] have been detected in the plasma of individuals with a diagnosis of schizophrenia [9, 11]. Furthermore, serum levels of IL-1β are elevated in the mothers of offspring who later develop psychosis [12]. Animal models of maternal immune activation (MIA) also provide evidence of elevated IFNγ and IL-1β levels in maternal serum, as well as the serum and brains of fetuses [6, 13]. Moreover, in mice exposed to MIA, offspring who are susceptible (those that show, as adults, deficits in social and cognitive functioning as seen in schizophrenia) have elevated plasma levels of IL-1β (among other cytokines), compared to control mice and mice that are resilient to MIA [14].

However, the impact and outcome following prenatal immune activation is heterogeneous between individuals [5, 14, 15]. For example, recent work in an MIA mouse model highlights the existence of subgroups of MIA-exposed offspring that show dissociable behavioural, transcriptional, neuroimaging, and immunological profiles [14, 16]. This is consistent with epidemiological studies in human cohorts: not all foetuses exposed to MIA will go on to develop schizophrenia [1, 17, 18]. It is therefore likely that MIA interacts with other factors, such as genetic background, to modulate the risk of developing schizophrenia. Indeed, the two-hit model of schizophrenia suggests that an amalgam of genetic risk and environmental insult is necessary to alter neurodevelopment enough to ultimately precipitate the symptoms of the disorder [19–21].

Importantly, if there are individual differences in human responses to immune activation due to genetic variability, there will certainly be differences in such responses between human and animal systems, given that they are even more genetically divergent. Although animal studies have provided important mechanistic insights, interactions between schizophrenia genetic burden and MIA cannot be fully recapitulated by rodent models: species differences in gene expression cannot be discounted [22–25]. It is therefore important to test the impact of gene-environment interactions in human model systems, such as human induced pluripotent stem cells (hiPSCs). These can be generated by reprogramming somatic cells, including hair keratinocytes and skin fibroblast biopsies, from patient cohorts or healthy controls [26–28]. The resulting hiPSCs can then be differentiated into multiple relevant cell types that retain the genetic make-up of the donor [29, 30] – including neural progenitor cells (NPCs), early precursors to neurons which are highly prevalent in the fetal brain [31]. There is evidence that hiPSC-derived NPCs closely resemble fetal brain tissue, recapitulating the neurodevelopmental hallmarks of the late first trimester/early second trimester stage [2, 29, 30, 32–34]. These facets make hiPSC-NPCs uniquely placed to model human fetal neurodevelopmental mechanisms and gene-environment interactions *in vitro* [33, 35].

Previous work from our group showed that transient IFNγ treatment of hiPSC-NPCs from healthy controls increases neurite outgrowth (a cellular phenotype associated with neurodevelopmental disorders) and disproportionately alters the expression of genes associated with schizophrenia and autism [2]. The aim of the current study is therefore to understand how specific cytokines (IFNγ and IL-1β) implicated in the association between MIA and schizophrenia risk influence transcriptional responses in early cortical neural progenitors. We use induced cortical neural progenitors with forebrain identity, as there is extensive evidence of prefrontal cortical abnormalities in patients with schizophrenia [7, 36, 37], and experiments in animal models have shown that exposure to inflammatory cytokines alters proliferation and differentiation of neural progenitors [10, 38]. We hypothesise that NPCs derived from patients with schizophrenia will respond differently to IFNγ and IL-1β compared to cells from healthy donors. If so, this may shed light on the mechanisms by which maternal immune activation increases the risk of developing schizophrenia.

## Materials and methods

### Participants

This study included hiPSC lines derived from six participants: three individuals with a diagnosis of schizophrenia (cell lines 138_SZM_09, 044_SCZ_04 and 115_SCZ_01) (Supplementary Figure 1) and three healthy donors with no history of psychiatric illness (cell lines M1_CTR_04, M2_CTR_42, M3_CTR_36S – previously described in [32, 33]). Participants were recruited as part of the Patient iPSCs for Neurodevelopmental Disorders (PiNDs) study (REC No 13/LO/1218). Participants with a diagnosis of schizophrenia were recruited at the Maudsley Hospital, London. The collection of data used for this research was approved by the NHS Research Ethics Committee at the South London and Maudsley (SLaM) NHS Research and Development Office. All participants gave written informed consent before contributing to the study. A diagnosis of schizophrenia was established based on International Classification of Diseases (10^th^ revision) [39] with the diagnosis (code F20), assessed using the Operational Criteria checklist [40] by a psychiatrist on the basis of information recorded by the clinical team following psychiatric interview. Healthy, unaffected individuals were selected as controls on the basis of having no history of psychiatric disorders [33].

### Reprogramming of keratinocytes

Hair root samples were collected by plucking occipital scalp hair (∼10+ roots per participant) and submerging these in Mouse Embryonic Fibroblast medium containing 50 µg/mL Gentamycin and 15 mM HEPES buffer (Gibco). The roots were then transferred to Geltrex™-coated 4-well plates (ThermoFisher), and outgrowth promoted, by supplementing with hair medium (Dulbecco’s Modified Eagle’s Medium (DMEM)

Advanced (Sigma Aldrich), GlutaMAX™ (ThermoFisher), 10% FBS (Clonetech), HEPES buffer and Gentamycin), to establish primary keratinocytes. The keratinocytes were subsequently reprogrammed into human induced pluripotent stem cell (hiPSC) lines. This transformation was induced by introducing Sendai viruses encoding Yamanaka Factors (human OCT4, SOX2, KLF4 and C-MYC), using a CytoTune-iPS 2.0 Sendai expressing Reprogramming Kit (ThermoFisher, A16517). The treated keratinocytes were plated onto an irradiated MEF feeder layer (Millipore) and supplemented Epilife medium. After ten days, Epilife medium was exchanged for hES medium, which was comprised of KO-DMEM/F12 supplemented with 20% knock-out serum, non-essential amino acids, Glutamax, β-mercaptoethanol (all from Life Technologies) and bFGF (10 ng/mL; Peprotech). After two more weeks, reprogrammed colonies were selected and plated on Nunc multi-plates (Thermo Scientific) coated with Geltrex (Life technologies) and supplemented with E8 media (Life Technologies).

### Maintenance of hiPSCs

The successfully reprogrammed hiPSCs were incubated in hypoxic conditions (5% CO2, 5% O_2_) at 37°C and maintained in StemFlex™ media (Gibco) on 6-well NUNC™ plates (ThermoFisher) coated with Geltrex™ (ThermoFisher). Cells were passaged (at a ratio between 1:6 and 1:18) upon reaching 60-70% confluency. During passage, cells were washed with room temperature Hank’s Balanced Salt Solution (HBSS) and incubated at 37°C with Versene (EDTA) solution (Lonza) for 3-5 minutes, then replated in new Geltrex™-coated NUNC™ plates.

### Directed differentiation of hiPSCs

The six hiPSC lines used in this study were then differentiated into forebrain cortical neural progenitor cells (NPCs) by dual SMAD inhibition [32, 33]. In preparation for neuralisation, hiPSCs were passaged onto 6-well NUNC™ plates coated with Geltrex™ at a 3:2 ratio and maintained under hypoxic conditions for ∼24–48 hrs until they approached 100% confluence. Directed differentiation was then initiated by changing StemFlex™ medium to neuralisation medium containing N2:B27 (N2 medium and B27 medium at a 1:1 ratio) supplemented with 100 nM LDN193189 (Sigma Aldrich) and 10 µM SB431542 (Sigma Aldrich) for dual SMAD inhibition. N2 medium consisted of DMEM/F12 (Dulbecco’s Modified Eagle’s Medium/Nutrient Mixture F12 Ham; Sigma Aldrich), supplemented with 1X GlutaMAX™ and 1X N2 supplement (ThermoFisher). B27 medium consisted of Neurobasal® medium (ThermoFisher), 1X GlutaMAX™ (ThermoFisher) and 1X B27 supplement without vitamin A (ThermoFisher).

The neuralised cells were then incubated under normoxic conditions (37°C, 5% CO_2_, 20% O_2_). Neuralisation medium was replenished every 24 hours from day 0 to day At the end of this 7-day neuralisation period, neuralisation medium was replaced with N2:B27 (without inhibitors), which was replenished every 24 hours from day 8 onwards. The neuralised cells were passaged four times: on day 7, day 12, day 15/16 and day 20/21. The passage procedure was, briefly, as follows: cells were washed with room temperature HBSS (ThermoFisher) and treated with Accutase (ThermoFisher) and incubated for 3–4 minutes at 37°C. The cells were then collected with the Accutase and mixed with room temperature DMEM/F12 (at a 2:1 ratio) and centrifuged at 1250 RPM for two minutes to separate the cells and Accutase. Cells were plated on new 6-well NUNC™ plates coated with Geltrex™. Passaging ratios were 1:1 for neural passaging 1 and 2, and 2:3 for neural passaging 3. To enhance cell survival, 10 µM protein kinase (ROCK) inhibitor (Sigma Aldrich), was added for 24 hours with the plating medium at each neural passage. After neural passage 3, cells were frozen in 10% DMSO (dimethyl sulfoxide). Cryovials were stored at -80°C for 24–48 hours in Mr. Frosty containers (to control freezing rate) before being transferred to liquid nitrogen.

For the final stages of neural passaging, cryovials were thawed in a 37°C water bath for 1 minute. The cell suspension was transferred to a 15 ml tube containing DMEM/F12 and centrifuged at 1250 RPM for 2 minutes. The cell pellet was resuspended in 3 mL of N2:B27 supplemented with 10 µM ROCK inhibitor and plated in Geltrex™-coated 6-well NUNC™ plates. From this point on, the following inhibitors were added to the NPC media (to make N2:B27-FGF): 10 ng/mL bFGF (basic Fibroblast Growth Factor; Peprotech), 100 μM β-mercaptoethanol (Life Technologies), 5 μg/mL insulin (Life Technologies), 1X non-essential amino acids (Life Technologies), 200 μM ascorbic acid (Sigma Aldrich). The cells were then expanded at a 1:3 ratio (in 2–5 neural passages) to prepare three wells of each line (one for each experimental condition).

Successful reprogramming of hiPSCs was validated as described in previous studies [32, 34, 41]. Pluripotency of all hiPSCs was confirmed by immunocytochemistry, differentiation of embryoid bodies into the three characteristic germ layers [42–45] (Supplemental Figure 1), and PluriTest analysis of Illumina HT12v4 transcriptome array data (https://www.pluritest.org) [46]. Alkaline phosphatase activity was further used to assess the pluripotency of hiPSCs using an alkaline phosphatase expression kit (Milipore). Genome integrity of hiPSC lines was assessed by an Illumina Human CytoSNP-12v2.1 beadchip array and analysed using KaryoStudio software (Illumina, San Diego, CA).

### Acute treatment with pro-inflammatory cytokines

NPCs were treated for ∼24 hours in three treatment conditions: IFNγ, IL-1β or vehicle. Media was fully removed and replaced with 3 mL per well of treatment media (N2:B27-FGF, supplemented as follows). IFNγ wells were treated with 25 ng/μL IFNγ (Abcam); the IL-1β wells with 10 ng/μL IL-1β (Abcam), as in [2]; and the control wells with vehicle (unsupplemented N2:B27-FGF media). After 24 hours, cells were lysed and collected in TRIzol® reagent (Thermo Fisher) and rapidly frozen on dry ice. The frozen samples were stored at -80°C until RNA extraction.

### RNA extraction and sequencing

RNA was extracted from the eighteen samples in two batches (to ensure durations of exposure of each sample to extraction reagents were well controlled). Both batches of extractions were conducted on the same day, by the same experimenter. The batches were randomised for experimental group (batch 1: lines M1_CTR, M2_CTR, 138_SZM; batch 2: lines M3_CTR, 044_SCZ, 115_SCZ), using the RNeasy Plus Mini Kit (QIAGEN), according to the manufacturer’s instructions. Extracted RNA was sent for sequencing at GENEWIZ® Ltd. Strand-specific, paired-end RNA sequencing with Poly(A) selection was performed using the Illumina® NovaSeq platform, at a read length of ∼30 million reads per sample.

### Quality control of RNA sequence and gene expression data

Initial quality control checks of raw RNA sequence data were conducted using the FastQC software from Babraham Bioinformatics (www.bioinformatics.babraham.ac.uk/projects/fastqc). Sequence reads were then aligned to the latest version of the human reference genome (hg38) using the STAR (Spliced Transcripts Alignment to a Reference) alignment tool [47]. The number of reads mapped onto each gene in Ensembl’s gene annotations for hg38 (version 99) was counted using FeatureCounts [48]. We plotted the distribution of log_10_-transformed counts-per-million (CPMs) and, by visual inspection, set a threshold of log_10_CPM = 0.6 for filtering out lowly expressed genes in order to minimise technical noise and reduce the multiple-testing burden (Supplementary Figure 7). After applying that threshold, 15060 out of 60642 genes were left for downstream analysis. TMM (trimmed mean of M-values)-normalization (Robinson & Oshlack, Genome Biology, 2010) was then applied on the gene counts, gene-expression values were log_2_-transformed, and observational-level theoretical variances were calculated using ‘voom’ for precision-weighting [49].

### Differential gene expression

In order to evaluate potential sources of overall gene expression variation, we performed Principal Component Analysis (PCA) on the *voom*-transformed gene expression [49], plotting the samples along the first three principal components (Supplementary Figure 2). We then used the ‘variancePartition’ R package [50] to estimate the contribution to the variance in expression of each gene for the following variables: study subject, clinical group (schizophrenia (SZ)/control), treatment condition (IFN-γ/IL-1β/vehicle), donor age, and the fraction of all RNA sequencing ‘reads’ that were mapped to genes (i.e., ‘assigned percent’).

To prepare the expression data for linear mixed effects modelling, we applied *voomWithDreamWeights* (‘variancePartition’ R package) to the expression-level-filtered TMM-normalized counts. We then conducted linear mixed model regressions using *dream* (‘variancePartition’), which allows modelling of interindividual variability by adding individual identifiers as a random effect in the regression model, as shown below [50, 51]. We also included the ‘assigned percent’ for each sample as a covariate, as we observed this variable to have a non-negligible effect on the overall variance of gene expression (Supplementary Figure 3). Participant age was not included as a covariate on the assumption that reprogramming samples to stem cells negates age-related effects [52, 53]. Age did seem to contribute to variation in gene expression in the variancePartition analysis (Supplementary Figure 3), but we attributed this to noise due to the small size of our sample. All the participants were male, so gender was not included as a covariate. The final model with an interaction term between clinical group and treatment was as follows:

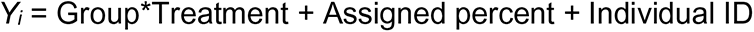

(Where “Group”, “Treatment” and “Assigned percent” were fixed effects and “Individual ID” was a random intercept effect).

Using contrasts, the following differential gene expression signatures were generated:

A. **Vehicle-treated SZ NPCs vs vehicle-treated control NPCs** (i.e., between SZ and control NPCs treated with vehicle).
B. **IFNγ-treated control NPCs vs vehicle-treated control NPCs** (i.e., the effect of IFNγ stimulation on gene expression in the control NPCs).
C. **IFNγ-treated SZ NPCs vs vehicle-treated SZ NPCs** (i.e., the effect of IFNγ stimulation on gene expression in the SZ NPCs).
D. **Interaction effect of IFNγ treatment in schizophrenia NPCs vs in control NPCs** (i.e., how the transcriptional response to IFNγ stimulation differs in SZ NPCs compared to control NPCs).
E. **IL-1β-treated control NPCs vs vehicle-treated control NPCs** (i.e., the effect of IL-1β stimulation on gene expression in the control NPCs).
F. **IL-1β-treated SZ NPCs vs vehicle-treated schizophrenia NPCs** (i.e., the effect of IL-1β stimulation on gene expression in the SZ NPCs).
G. **Interaction effect of IL-1β treatment in SZ NPCs vs in control NPCs** (i.e., how the transcriptional response to IL-1β stimulation differs in SZ NPCs compared to control NPCs).

Approximation of residual degrees of freedom and subsequent calculation of moderated eBayes *t*-statistics was done using the Satterthwaite method in ‘dream’ [51].

### Gene set enrichment analysis

Our gene set enrichment analyses (GSEA) included 935 unique gene sets: 519 immune-related and 421 nervous-system/neural function related (5 overlapping). Of these, 135 were obtained from previous literature [54–57] and the remaining from either the Molecular Signature Database (HALLMARK and Gene Ontology biological process gene sets) or the pathway databases KEGG, PANTHER, Pathway Commons and Reactome (see Supplementary Tables 4A-G for a full list of gene sets). GSEA assesses whether genes belonging to specific pathways or predefined sets of genes are over-represented in the significant or peri-significant results of a differential expression analysis. We used a linear mixed effects regression-based competitive gene set enrichment approach using the GSEA tool (Subramanian et al, 2005). GSEA was run on the seven signatures generated by the DGE analysis. We applied the *fgsea* function of the R package ‘fgsea’ [58], using the standardized Z-score obtained in the differential expression analysis to rank the genes and running 100,000 permutations. All gene sets containing fewer than five genes were excluded. Multiple testing correction was performed within fgsea using the false discovery rate (FDR) method, and gene sets with an FDR < 0.05 were considered significant.

The resulting gene sets showed substantial constituent similarity (Supplementary Figure 4), so we clustered them based on the overlap of the genes that belong to each gene set. This was done by calculating the Jaccard Similarity Index (which quantifies the intersection of two lists) between all pairs of significantly enriched gene sets and then applying a hierarchical clustering of gene sets based on the resulting dissimilarity matrix (1-Jaccard similarity). We then applied a cut-off of *h=*0.5 to the dendrograms to obtain clusters of significantly enriched gene sets (Supplementary Figure 5).

### Enrichment of schizophrenia genes

To test whether differentially expressed genes in our experimental conditions were enriched for genes differentially expressed in *post-mortem* brain samples originating from SZ cases, we split genes by direction of effect (up- or downregulation) and assessed their overlap with genes differentially expressed in schizophrenia, according to Gandal et al [59]. Significance of this overlap was estimated using the Fisher’s exact test in R, through the *GeneOverlap* package, assuming a genome size of 20000 protein-coding genes.

To test whether the differentially expressed genes in our experiments were enriched for GWAS-supported genes, we performed gene-set enrichment analysis using MAGMA. The summary statistics file from the schizophrenia GWAS performed by Pardiñas et al [60] was downloaded and pre-processed using standard quality control procedures, where variants with minor allele frequency < 0.01, or those in the extended MHC region on chromosome 6, from 25-34 Mb, were removed. The GWAS variants were annotated to a list of protein-coding genes provided by the authors, which included genes located on chromosomes 1 to 22 and X, allowing a window of 35 kb upstream and 10 kb downstream of each gene, as described previously [61]. Gene-level enrichment analysis was performed to identify genes more likely to be associated with schizophrenia according to the GWAS results, using the European subset of the 1000 Genomes Phase 3 as reference panel. Subsequently, we tested whether there was an enrichment of genes differentially expressed in our experimental models (excluding those that were non-coding) within these results. All tests were corrected for multiple testing using the false discovery rate method, according to the number of gene sets analysed per condition (i.e., two gene lists [up- and downregulated genes] from two groups each [control vs. SZ cell lines] = 4 comparisons per analysis).

## Results

### Demographic and sample details

The demographic and clinical characteristics of the six participants are described in Table 1. Subjects were male and of White British, or ‘Other White’ background. Ages ranged from 33 to 55 years old. The patients were diagnosed with paranoid schizophrenia, and controls were selected on the basis of having no history of neuropsychiatric disorders.

**Table 1.** Demographic and sample details.

| Cell line | Diagnosis | Year diagnosed | Medication | Age | Gender | Ethnicity | Reprogrammed by |
| --- | --- | --- | --- | --- | --- | --- | --- |
| 044 | Schizophrenia | 2011 | Risperidone | 33 | Male | White British | Sendai virus |
| 115 | Schizophrenia | 2010 | Aripiprazol | 43 | Male | White British | Sendai virus |
| 138 | Schizophrenia | 2008 | Risperidone,<br>Mirtazapine | 39 | Male | Black British | Sendai virus |
| M1 | Control | - | - | 55 | Male | White British | Lentivirus |
| M2 | Control | - | - | 35 | Male | White British | Lentivirus |
| M3 | Control | - | - | 35 | Male | White British | Sendai virus |

### Validation of hiPSCs and NPCs

All hiPSC lines differentiated into embryoid bodies with three characteristic germ layers and expression of pluripotency markers NANOG, OCT4, SSEA4 and TRA-1-81 (Supplementary Figure 1). For each participant, one clone was used for the NPC induction. All hiPSC lines successfully differentiated into NPCs as determined by immunostaining for known NPC markers βIII-tubulin and Nestin (Figure 1).

**Figure 1.**
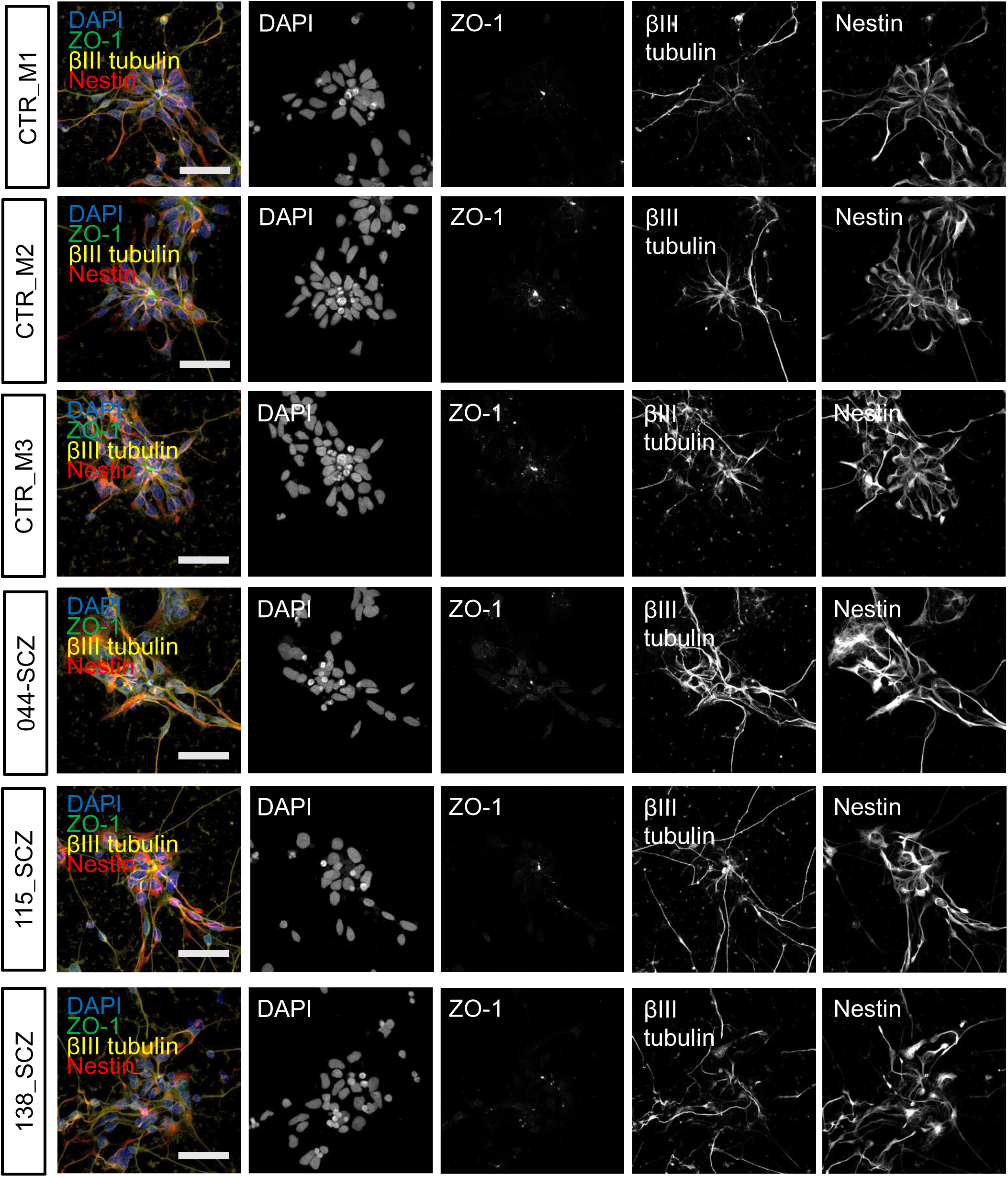
*Validation of neural progenitor cells.* Successful differentiation to neural progenitor state was confirmed by staining at Day 20 for NPC markers, Nestin and β-III-tubulin. DAPI was used for baseline nuclear staining. Scale bar = 50µm

### Sources of variation in gene expression

We observed that the greatest source of variability across all samples was individual differences between the patients, as seen with principal component (PC) 1 (Supplementary Figure 2 A-C, left panels), as well as in the VariancePartition (Supplementary Figure 3) comparison. The VariancePartition analysis also revealed that the other variables included in the model (while having a low impact on overall gene expression variation individually) together explained at least 50% of the expression variance (Supplementary Figure 3). The importance of the clinical group was further supported by the clustering of SZ and control samples along the 2nd and 3rd principal components of PCA analysis (Supplementary Figure 2C, right panel).

### Differential expression of genes and gene set enrichment analysis (GSEA)

Of the seven comparisons we made, four showed significant differential gene expression at *FD*R < 0.05: the effect of diagnosis (Signature A), and the three IFNγ treatment conditions (Signatures B-D). There were no statistically significant (*FDR* 5%) differentially expressed genes (DEGs) as a result of acute IL1-β treatment. A full table of DEGs can be found in Supplementary Tables 3A-G. Immune-related gene sets were among the top ten most significantly enriched for all seven signatures, and synaptic transmission-related gene sets were among the top ten in four of the seven signatures. A full table of enrichment terms can be found in Supplementary Tables 4A-G. Details of DEGs and gene set enrichment analysis (GSEA) results for each signature are presented below.

### Effect of schizophrenia diagnosis (Signature A)

We first investigated the gene expression differences observed in SZ relative to control lines, in vehicle-treated NPCs. We found only one statistically significant DEG (Figure 2A), *AL132709.7*, a human-specific lncRNA gene which was overexpressed in patient lines (*FDR =* 0.0395; logFC = 3.111). Our sample was underpowered to detect other DGEs, but the GSEA revealed 26 significantly enriched gene-sets. The top five gene sets were enriched among nominally significant upregulated genes in this comparison (Figure 2B), and the gene set with the lowest *p-*value was ‘Lek2015 loss-of-function (90)’ (*FDR* = 0.00098; normalized enrichment score (NES) = 1.36; genes in gene set = 3007), which contains 3007 genes that are intolerant to loss-of-function variants. This is consistent with previous schizophrenia genetic association studies that find associations with the same loss-of-function gene sets [55, 56].

**Figure 2.**
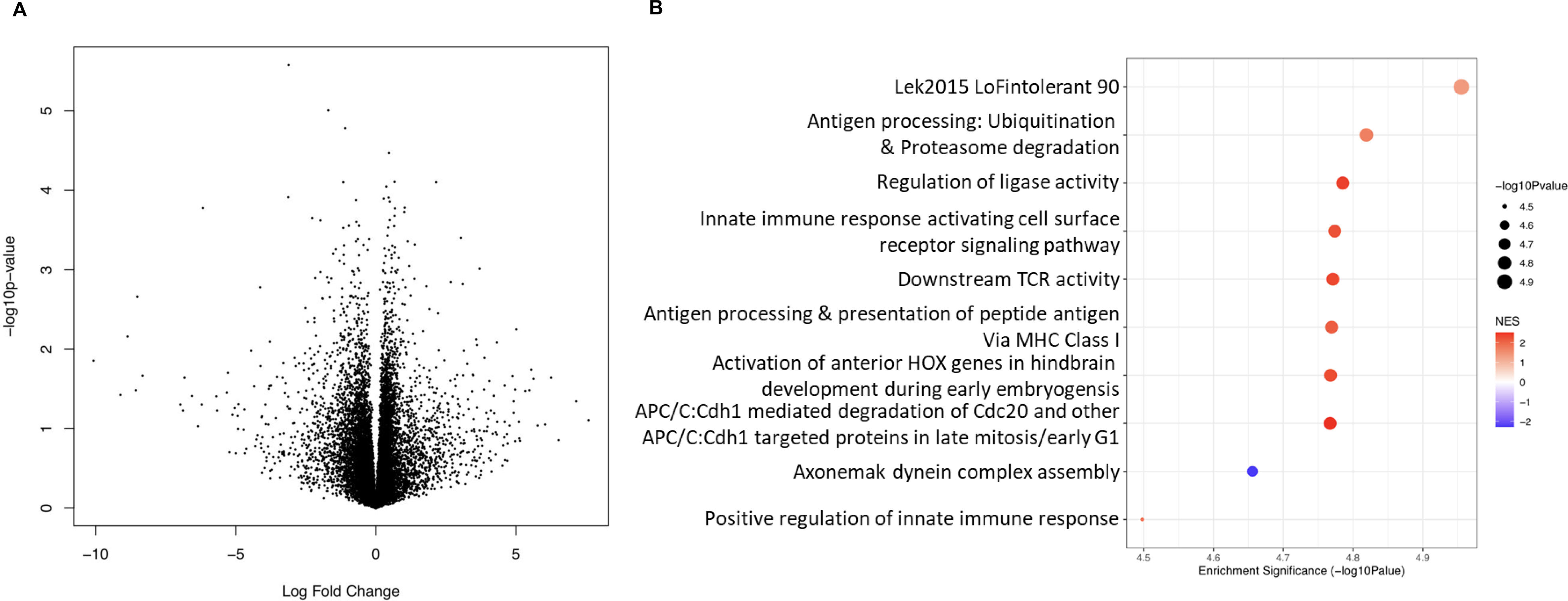
Expression differences between NPCs from cases vs. controls (signature A) at the gene and pathway level. **A.** The y-axis here shows statistical significance (-log_10_ p-value) of differential expression of genes in untreated cells from patient donors compared to gene expression in cells from untreated control donors. The x-axis shows the log_2_ fold change of expression of those genes in schizophrenia cell lines vs control cell lines. **B.** The top 10 significantly enriched gene set clusters (the gene set with the lowest p-value in each cluster is labelled on the x-axis). Data-points are sized according to significance (-log_10_ p-value) and coloured according to normalised enrichment score (NES), with blue indicating downregulation and red indicating upregulation.

### Effect of IFNγ treatment in control NPCs (Signature B)

We observed 1847 upregulated and 1533 downregulated genes in control NPCs (total = 3380 genes, out of 15061 tested) in response to the IFNγ treatment, relative to the vehicle-treated lines (Figure 3A and Supplementary Table 1). We observed significant upregulation of *STAT1* (*FDR =* 5.572 x 10^-6^; logFC *=* 5.680)*, STAT2* (*FDR =* 5.045 x 10^-6^; logFC *=* 2.247) and *JAK2* (*FDR =* 0.001; logFC = 2.111) – consistent with activation of the IFNγ signal transduction pathways) – as well as *IRF1* (*FDR =* 3.23 x 10^-6^; logFC *=* 7.022), a key downstream signalling target of this cytokine [2, 62]. The genes whose expression was most significantly altered by IFNγ treatment were *IFI27* (*FDR* = 2.97 x 10^-6^; logFC = 6.067) and *CD274* (*FDR* = 2.97 x 10^-6^; logFC = 6.386), both upregulated. Three genes encoding guanylate-binding proteins were among those that showed the highest fold change (logFC): *GBP1* (*FDR* = 3.11 x 10^-5^; logFC = 14.622) *GBP5* (*FDR* = 0.0037; logFC = 13.473) and *GBP4* (*FDR* = 0.0002; logFC = 13.113). The upregulation of these genes is consistent with the role of guanylate-binding proteins (especially *GBP1*) in the inflammatory response associated with IFNγ [63]. Our findings for this comparison were also consistent with recent work which found MHC-I related genes among the most differentially expressed in IFNγ-treated control neural progenitors and neurons [2]. In our results (Supplementary Table 3A), key MHC-I related genes such as *HLA-A, HLA-B* and *HLA-C* are all consistently upregulated in response to IFNγ treatment.

**Figure 3.**
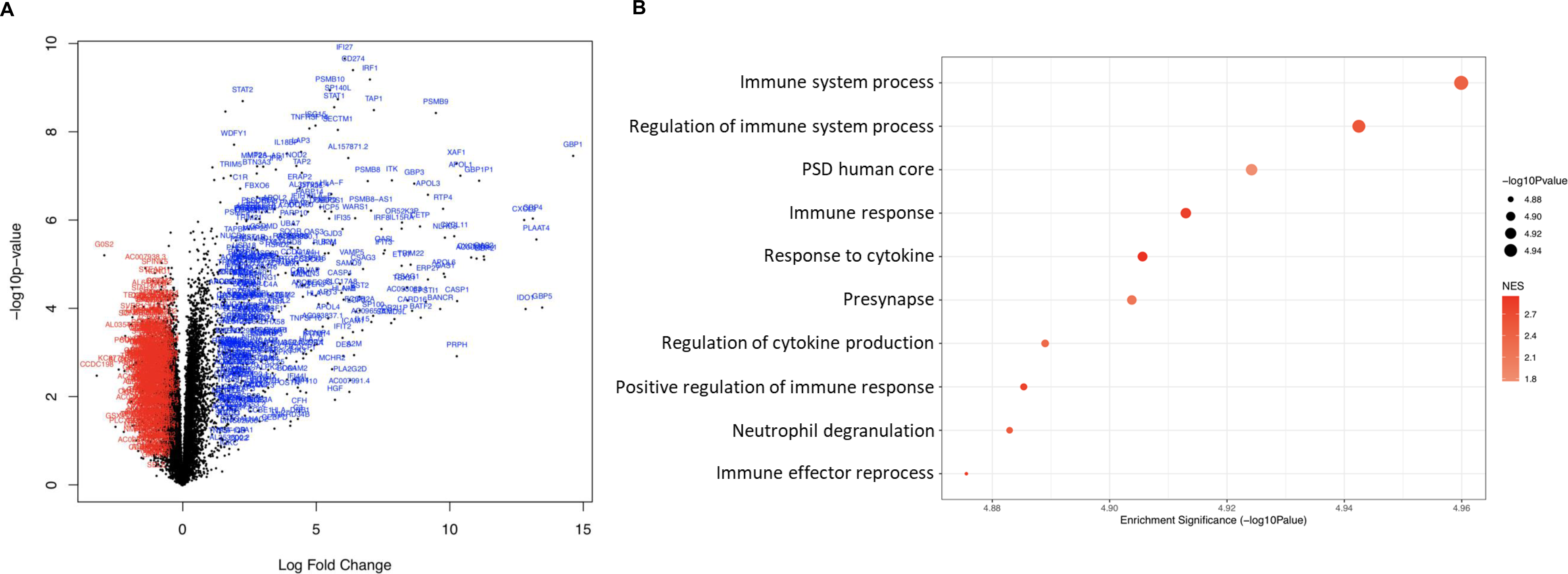
Expression differences between NPCs from IFNγ treated versus untreated control NPCs (signature B) at the gene and pathway level. **A.** The volcano plot shows, on the y-axis, the statistical significance (-log_10_ p-value) of differential expression of genes in IFNγ-treated control NPCs compared to untreated control NPCs. The x-axis is the magnitude of change (log_2_ fold change) in expression of those genes due to after IFNγ treatment. **B.** The top 10 significantly enriched gene set clusters (the gene set with the lowest *p*-value in each cluster is labelled on the y-axis). Please see Supplementary Spreadsheets 4A-G for full lists of enriched gene sets for each of the signatures. Data-points are sized according to significance (-log_10_ p-value) and coloured according to normalised enrichment score (NES), with blue indicating downregulation and red indicating upregulation.

We additionally tested whether the genes differentially expressed due to IFNγ treatment overlapped with genes known to be differentially expressed in patients with SZ, based on gene lists provided by [64] (split by direction of effect: up- or down-regulation) and using the R package *GeneOverlap* [65]). We observed a significant overlap between the genes downregulated in the IFNγ-treated control NPCs with genes downregulated in SZ (p = 7.00 x 10^-4^, *FDR* = 0.0014, odds ratio (OR) = 1.3), but not with those that were upregulated (p < 0.05). Similarly, the genes upregulated in our model overlapped with those upregulated in SZ (p = 1.20 x 10^-8^, *FDR* = 4.80 x 10^-8^, OR = 1.5), but not with those that were downregulated (p < 0.05). We performed another gene-set enrichment analysis, using MAGMA [61], to test whether DEGs in our model were overrepresented in GWAS summary statistics from a large-scale schizophrenia GWAS [55]. This analysis did not identify evidence that IFNγ-regulated genes in our model were associated with interindividual genetic variation contributing to schizophrenia susceptibility (p < 0.05). Collectively, these results suggest that IFNγ signalling may impact neurodevelopment in a way that predisposes to schizophrenia, but that this may be independent from genetic effects. One caveat of the enrichment analysis using MAGMA is that it excludes genes within the MHC region due to the complex linkage disequilibrium structure at this locus, even though many of the genes in the MHC region are relevant for the IFNγ response.

We further observed 168 pathways enriched when comparing IFNγ-treated cells and untreated cells in control NPCs. The gene set with the lowest *p-*value (Figure 3B) was ‘immune system process’ from Gene Ontology (*FDR* = 0.0002; *NES* = 2.41; genes in gene set = 1235), which consists of genes involved in the development or functioning of the immune system. All of the top ten gene pathways for this signature were overexpressed amongst genes upregulated in response to IFNγ treatment. While most of these were related to the immune response initiated by cytokine exposure, we observed two that were, notably related to synaptic function: ‘post-synaptic density, human core’ and ‘presynapse’.

### Effect of IFNγ treatment in schizophrenia NPCs (Signature C)

We observed 1061 upregulated and 919 downregulated genes in SZ cell lines (*FDR <* 0.05, total = 1980 genes, out of 15061 tested) in response to the IFNγ treatment, relative to the vehicle-treated SZ lines (Figure 4A and Supplementary Table 2). The genes whose expression was most significantly altered by IFNγ treatment in the SZ neural progenitors were *STAT2* (*FDR* = 1.46 x 10^-5^; logFC = 2.6103), *IFI27* (*FDR* = 1.74 x 10^-5^; logFC = 6.331) and *STAT1* (*FDR* = 1.74 x 10^-5^; logFC = 5.453). Once again, *IRF1* (*FDR* = 3.543 x 10^-5^; logFC = 7.278) and *JAK2* (*FDR* = 0.0033; logFC = 1.846) were also significantly upregulated. Here too, the highest logFC was shown by *GBP1* (*FDR* = 3.91 x 10^-5^; logFC = 12.308), followed by the pseudogene *GBP1P1* (*FDR* = 0.0001; logFC = 11.116).

**Figure 4.**
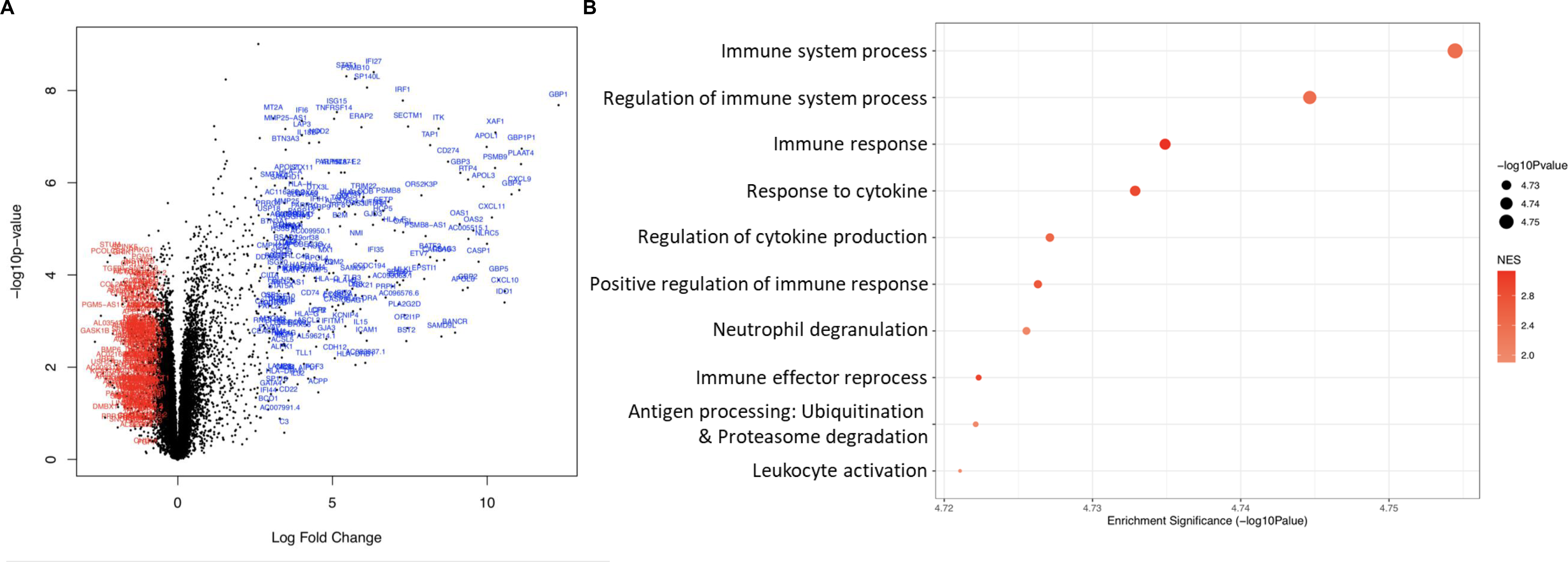
Expression differences between NPCs from IFNγ treated versus untreated schizophrenia NPCs (signature C) at the gene and pathway level. **A.** The volcano plot shows, on the y-axis, the statistical significance (-log_10_ p-value) of differential expression of genes in IFNγ-treated schizophrenia (SCZ) NPCs compared to untreated SCZ NPCs. The x-axis is the magnitude of change (log_2_ fold change) in expression of those genes due to after IFNγ treatment. **B.** The top 10 significantly enriched gene set clusters (the gene set with the lowest *p*-value in each cluster is labelled on the y-axis). Please see Supplementary Spreadsheets 4A-G for full lists of enriched gene sets for each of the signatures. Data-points are sized according to significance (-log_10_ p-value) and coloured according to normalised enrichment score (NES), with blue indicating downregulation and red indicating upregulation.

We saw that genes upregulated in SZ NPCs treated with IFNγ (*FDR* < 0.05) also significantly overlapped with those known to be upregulated in SZ patients (p = 1.10 x 10^-14^, FDR = 4.40 x 10^-14^, odds ratio (OR) = 1.9) (but not those that were downregulated). However, the downregulated genes in the model did not overlap with those downregulated (or upregulated) in cases, suggesting that the SZ NPCs may respond differently to IFNγ when compared to NPCs from unaffected individuals. Ultimately, we did not observe an enrichment of GWAS-supported variants within the genes differentially expressed in SZ NPCs after treatment, as observed in control NPCs (i.e., Signature B).

Nevertheless, there were 132 pathways enriched in the comparison of IFNγ- treated SZ cells (versus untreated SZ lines); the gene set with the lowest *p-*value (Figure 4B) was, again, ‘immune system process’ (*FDR* = 0.0002; *NES* = 2.366; genes in gene set = 1235). The results in this comparison show activation of similar pathways in response to IFNγ in SZ lines as seen in control lines in the previous comparison (Figure 4B). However, the transcriptional response appears attenuated, as we observed fewer DEGs overall.

### Interaction effect of IFNγ treatment and schizophrenia diagnosis (Signature D)

To get a general picture of whether the patient NPCs respond differently to IFNγ treatment compared to how control NPCs do, we first examined the overlap of DEGs between signatures B and C (presented as a Venn diagram in Supplementary Figure 6). It was evident that of the 4137 genes that responded to IFNγ in any of the two groups, only 1223 genes were in common to both, meaning that there are 2914 genes that appeared to respond differentially to IFNγ treatment between SZ and control NPCs. The following signature (Signature D) effectively assesses the same overlap; but subjects this comparison to an additional test of statistical significance (essentially omitting any of the 2914 genes observed in the initial comparison that may have differentially responded to treatment by chance). For this interaction term, we performed multiple-testing correction on the *p*-value obtained for the 4137 genes that are differentially expressed in response to IFNγ in any condition (controls and/or SZ cells). At *FDR* < 0.05 there were 359 genes that respond significantly differently to IFNγ between control and SZ cells (Figure 5A; Table 2); most significantly the mitochondrial complex genes *NDUFA2* (*FDR* = 0.0003; logFC = -0.591) and *NDUFS3* (*FDR* = 0.0006; logFC= -0.330) and the lncRNA gene *AC092279.2* (*FDR* = 0.0006; logFC = 0.645). Indeed, the previous comparisons show that the mitochondrial genes are overexpressed in response to IFNγ in the control NPCs (logFC = 0.405, *NDUFA2*; logFC = 0.145, *NDUFS3*) but are underexpressed in response to IFNγ in SZ NPCs (logFC = -0.187*, NDUFA2*; logFC = -0.185, *NDUFS3*). *AC092279.2* shows the opposite profile, responding with underexpression in control cells (logFC = - 0.517) and overexpression in SZ cells (logFC = 0.123).

**Figure 5.**
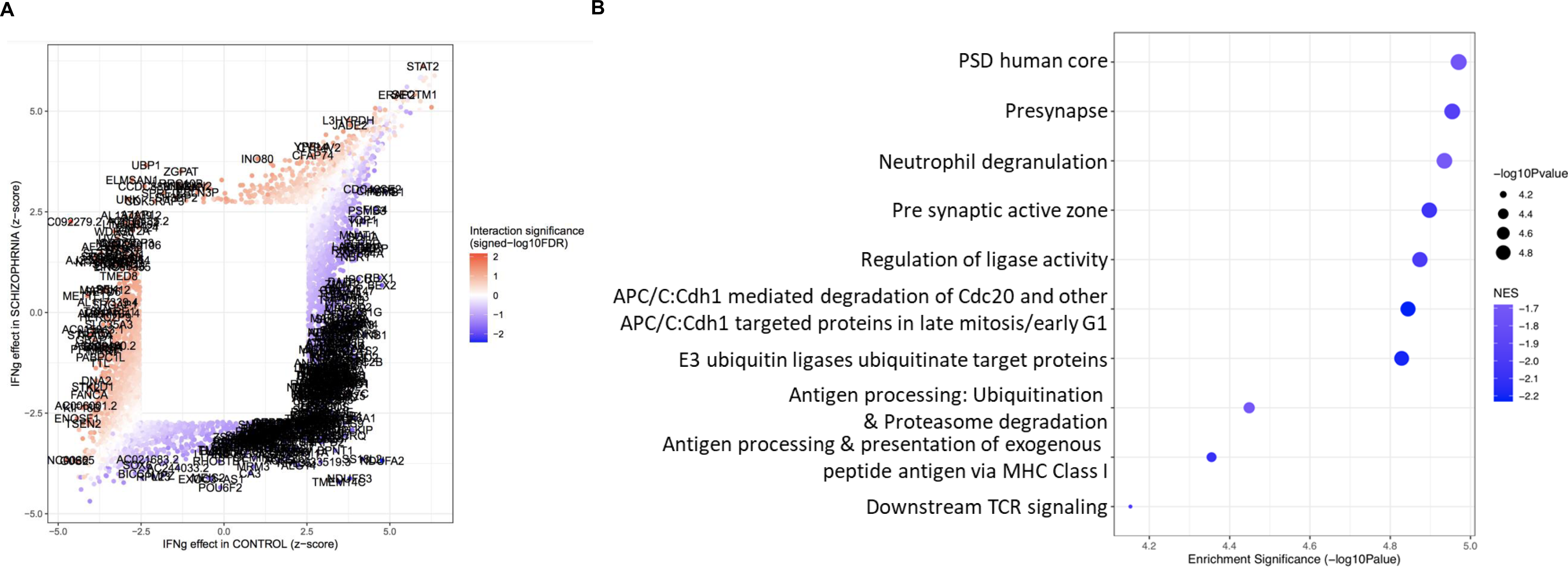
Interaction effect between IFNγ-treatment and diagnostic group on gene expression (Signature D). **A.** The scatterplot shows IFNγ response results for Signature in the 4137 genes that responded differentially to IFNγ in Signatures B and C. DEGs for control cells are on the *x*-axis and DEGs for schizophrenia cells are on the *y*-axis. The data are coloured by signed -log_10_FDR obtained for the interaction term (with blue indicating downregulation and red indicating upregulation). The 359 significant genes that are significant in the interaction are labelled. **B.** The top ten significantly enriched gene set clusters (the gene set with the lowest *p*-value in each cluster is labelled on the y-axis). Data-points are sized according to significance (-log_10_ p-value) and coloured according to normalised enrichment score (NES), with darker blue indicating greater downregulation.

**Table 2.**
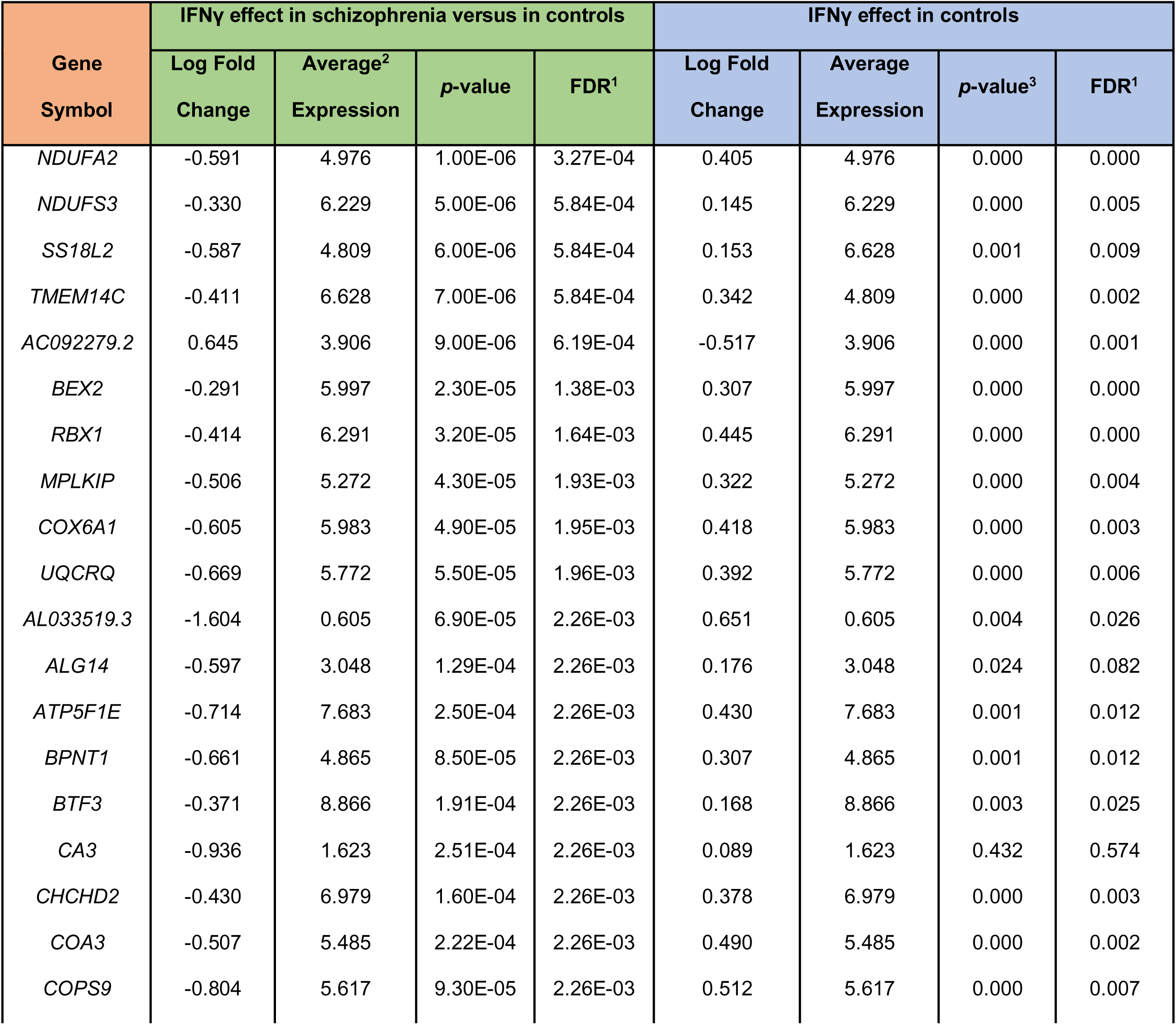
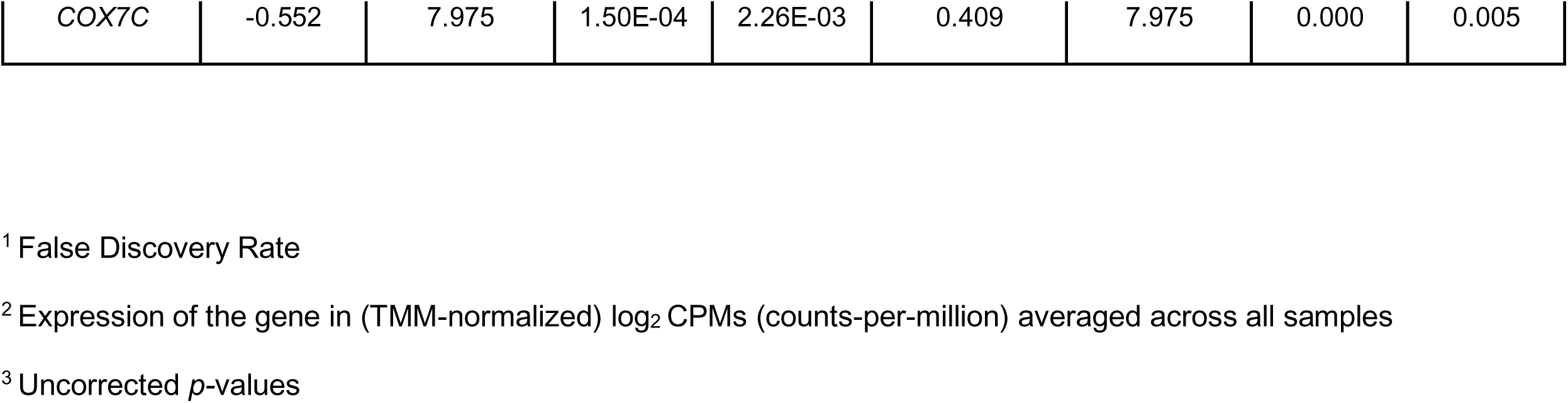
Top 20 genes significantly differentially expressed in IFNγ-treated compared to untreated cell lines in schizophrenia versus controls (Signature D) – all of these are downregulated. The right side of the table shows the effect of IFNγ treatment on the same genes in controls only, for comparison. Please see Supplementary Spreadsheet 3D for differential expression results for all genes in this comparison. A negative logFC indicates downregulation.

**Table 3.**
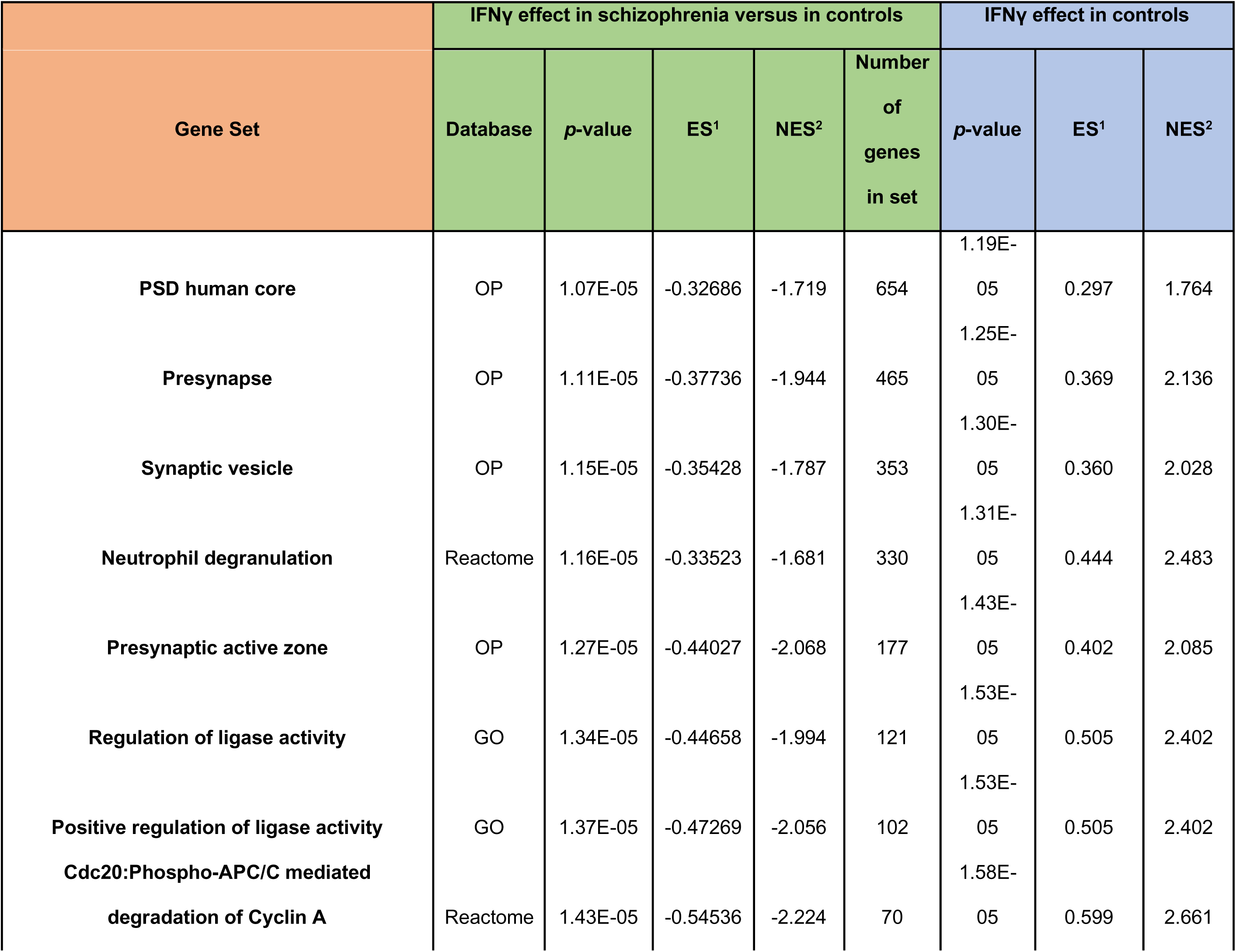
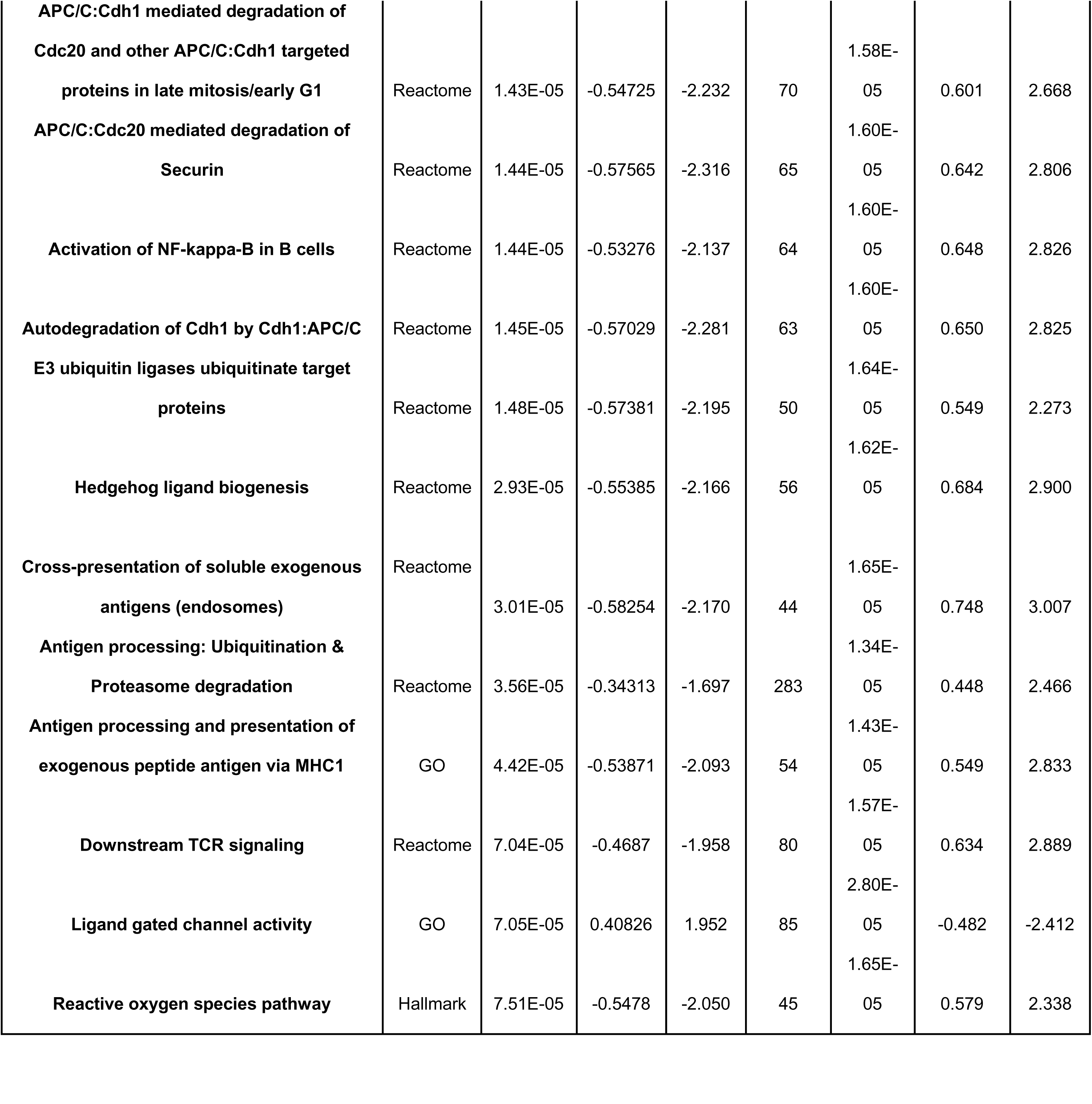
Top 20 gene sets significantly overrepresented among DEGs in IFNγ-treated compared to untreated cell lines in schizophrenia versus controls (Signature D). The right side of the table shows the effect of IFNγ treatment on the same genes in controls only, for comparison. Please see Supplementary Spreadsheet 4D for differential expression results for all genes in this comparison.

There were 20 gene sets that were differentially expressed in this comparison. These gene sets were comprised of genes that showed different transcriptional responses to IFNγ in SZ NPCs compared to control NPCs (Figure 5B). The most significantly different of these gene sets were ‘post-synaptic density (PSD), human core’ (*FDR* =0.001; *NES* = -1.72; genes in gene set = 654), which includes several notable genes including the Alzheimer’s risk gene *APOE*, autism and schizophrenia risk genes *NRXN1, CYFIP1* and *SHANK1-3*, NMDA receptor gene *GRIN1,* and *DLG4,* which encodes the postsynaptic density protein PSD-95; as well as a ‘presynapse’ gene set, which includes genes that regulate the pre-synaptic ‘active zone’ and synaptic vesicle formation [54, 55, 66] – notable genes in this gene set include *SV2A, MAOA* and several Na+/K+ transport ATPase genes. In other words, genes influencing synaptic transmission showed a particularly attenuated response to IFNγ treatment in SZ NPCs.

### Effect of IL-1β treatment on gene expression (Signature E, F & G)

We observed no differentially expressed genes associated with the effect of IL-1β treatment in either control (Signature E, Supplementary Figure 8A) or SZ (Signature F, Supplementary Figure 9A) NPCs (FDR > 0.05). We hypothesized that this could have been caused by the reduced expression of the main IL-1β receptors in cells at the neural progenitor stage. We assessed the expression of the genes encoding the IL-1β receptors *IL1R1, IL1R2,* and *IL1RAP,* and compared these to the expression of the IFNγ receptor genes *IFNGR1* and *IFNGR2*. We observed that the IL-1β receptor genes exhibited visibly lower expression relative to the IFNγ receptors (Figure 6), which may explain the lower responsiveness of the NPCs to IL-1β stimulation. In addition, these results suggest that IFNγ signalling may be more relevant than IL-1β at this developmental stage and/or for this cell type (NPCs).

**Figure 6.**
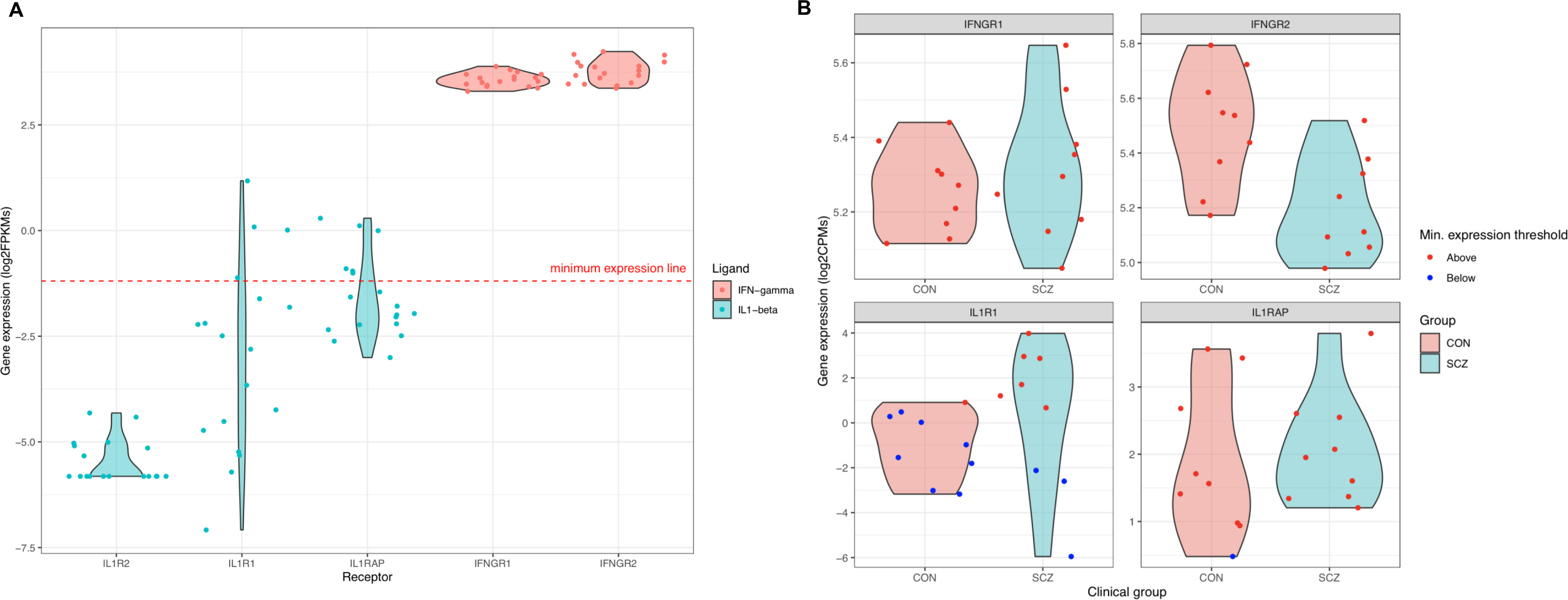
Expression of cytokine receptors in schizophrenia cell lines compared to controls. **A.** Shows the differential gene expression in schizophrenia (SCZ) compared to controls (CON) the four cytokine receptor genes that passed low-expression filtering. Values that lie above minimum expression threshold (log_2_ CPM = 0.6) are shown in red. **B.** Shows the differential gene expression in schizophrenia (SCZ) compared to controls (CON) the four cytokine receptor genes that passed low-expression filtering. Values that lie above minimum expression threshold (log_2_ CPM = 0.6) are shown in red.

Nevertheless, we observed 123 and 112 gene sets enriched for signatures E and F, respectively (Supplementary Figure 8B and 9B). For signature E, ‘regulation of immune system process’ was the most significant term (*FDR* = 0.0005; *NES* = 1.55; genes in gene set = 888); and ‘Lek2015 loss-of-function (90)’ [54, 55, 66] was the most significant term for signature F (*FDR* = 0.0008; *NES* = -1.97; genes in gene set = 3007). The fact that there are significant gene set enrichment terms for this comparison despite there being no DEGs suggests that there are indeed effects of IL-1β on transcription, but that our sample is underpowered to detect these individually. This method incorporates the expression signal from peri-significant genes and restricts the number of tests performed, reducing the multiple testing burden (instead of analysing 15061 genes for differential expression, as in the DGE analysis, our enrichment analysis tests 895 gene sets).

As there were gene sets significantly enriched for these signatures, we also tested whether genes that were differentially expressed at a more lenient threshold of *FDR* < 0.1 were enriched for genes differentially expressed in SZ. We observed that genes upregulated in SZ NPCs treated with IL-1β showed a significant overlap with those upregulated in schizophrenia cases (p = 0.0013, FDR = 0.0052, odds ratio (OR) = 1.6). Further, using MAGMA, we observed that genes downregulated in the SZ NPCs were enriched with GWAS-supported variants associated with SZ (β = 0.19, SE = 0.06, P = 8.56 x 10^-4^, FDR = 3.42 x 10^-3^). While these results corroborate a role for IL-1β signalling in schizophrenia, it is likely that the effects of this cytokine on NPCs are limited due to the lack of other cell types (such as microglia) in the culture system used or occur at another this involves another developmental stage.

The interaction effect between IL-1β and SZ (Signature G) also did not yield any significantly differentially expressed genes (Supplementary Figure 10A) but did yield 15 gene sets that were significantly enriched. Of these, ‘regulation of ligase activity’ had the lowest *p-*value for signature G (*FDR* =0.0015; *NES* = -2.15; genes in gene set = 121; Supplementary Figure 10B). These enrichment results also show a suppression of transcriptional response to IL-1β exposure in SZ NPCs, with pathways regulating the post-synaptic density and presynapse once again amongst the top ten (Supplementary Figure 10B).

## Discussion

In this study, we sought to assess how the cytokines interferon-gamma (IFNγ) and interleukin-1 beta (IL-1β) interact with genetic profiles associated with schizophrenia, to better understand the increased susceptibility to schizophrenia seen in offspring of mothers exposed to infection during pregnancy. We hypothesised that cortical neural progenitor cells (NPCs) derived from patients with schizophrenia would respond differently to IFNγ or IL-1β exposure compared to those of healthy controls.

We performed a preliminary analysis to evaluate the transcriptomic differences between our schizophrenia and controls NPCs without treatment stimulation and found only one differentially expressed gene. This result contrasts with the at least 80 DEGs found in schizophrenia patient-derived NPCs by Evgrafov et al [66]. However, the fact that our sample size is small and that we use a more stringent significance threshold (*FDR* < 0.05 as opposed to *FDR* < 0.1) may account for that difference. The identification of 26 gene-sets significantly enriched for this comparison further supports the idea that, while there are noticeable differences between schizophrenia and control NPCs at the transcriptomic level, our study may be underpowered to detect those for each gene individually.

We then assessed whether IFNγ treatment alters transcriptional responses in control neural progenitors and found that there were 3380 significant differentially expressed genes in response to IFNγ treatment. This is interesting itself, as the cell cultures used in this study do not contain glial cells – supporting the notion presented by our group in a recent study [2] that human NPCs can launch an immune response independent of microglia. Immune responses in the brain are thought to be predominantly mediated by glia [67], but the fact that NPCs are themselves responsive to a proliferation of IFNγ indicates that immunity in the brain extends beyond glial cells. This is also consistent with [68], who show that neuronal co-culture with activated microglia is sufficient to induce deficits in the neurons – suggesting that cytokines might come from glia, but neurons can respond to them independently. Moreover, in both control and schizophrenia NPCs, IFNγ treatment activated the canonical *JAK-STAT* signalling pathway, as would typically be seen in response to viral infection. Our findings for this comparison were also consistent with recent work which found MHC-I related genes among the most differentially expressed in IFNγ-treated control neural progenitors and neurons [2]. In our results (Supplementary Table 3A), key MHC-I related genes such *as HLA-A, HLA-B* and *HLA-C* are all consistently upregulated in response to IFNγ treatment. However, the genes most significantly upregulated on IFNγ exposure were *IFI27* and *CD274*. *IFI27* encodes Interferon Alpha Inducible Protein 27, which is involved in interferon-induced apoptosis and is considered to be a biomarker that differentiates between viral and bacterial infection [69]. *CD274* encodes a receptor ligand that binds to PD-1 receptors on T-cell surfaces, inhibiting T-cell activation and antibody production – an essential process for preventing autoimmunity [70]. Interestingly, there is some evidence that *IFI27* is differentially expressed in transgenic mice that exhibit schizophrenia-like behaviours [71]. Similarly, *CD274* is a member of several gene-sets found to be enriched in a study of *de novo* copy number variant associated to schizophrenia risk [72]. In light of this evidence, these genes may be promising candidates for future studies exploring the link between MIA and schizophrenia risk.

In schizophrenia NPCs, there were fewer differentially expressed genes in response to IFNγ treatment: only 1980. This may be because these cells are not able to activate in response to IFNγ to the same extent as controls. Physiologically, this could mean that cells from healthy donors exhibit a compensatory transcriptional response to infection, which is inhibited in cells from donors with schizophrenia. The two genes showing the most divergent response to IFNγ in SCZ and control cells (*NDUFA2* and *NDUFS3*) being mitochondrial complex I genes further suggests that schizophrenia donor cells are relatively driven to conserve energy in response to an infection, while healthy donor cells are able to expend more energy to restore health [68]. Indeed, there is some recent evidence for differences in expression of mitochondrial genes in rodents susceptible to MIA [14]. This may also be related to pre-existing deficits in mitochondrial function in schizophrenia lines, as it is well established that mitochondrial dysfunction contributes to the pathophysiology of schizophrenia [73].

The gene sets significantly enriched for IFNγ treatment in both schizophrenia and control NPCs largely converged in function, as expected, upon immune regulation. The gene sets that responded most differently to IFNγ in schizophrenia lines were those regulating the postsynaptic density, presynapse, and presynaptic active zone. This result shows that the genes involved in the aberrant response to immune activation by schizophrenia NPCs are involved in synaptic transmission, which fits with previous work by Shatz [82], as well as models of schizophrenia from other fields of neuroscience, including the concept of schizophrenia as a disorder of synaptic ‘dysconnection’ in computational neuroscience – a promising bridge between two very different but equally rich views of the same disorder. The dysconnection hypothesis suggests a dysregulation of neuromodulation (particularly across glutamatergic synapses) lies at the core of the various factors contributing to schizophrenia susceptibility [74, 75].

Our results did not reveal any significant DEGs in response to IL-1β treatment in SCZ or control NPCs. This is likely due to the low expression of the IL1 receptor 1 (IL1R1) gene that we observed, as blocking IL1R1 has previously been shown to significantly reduce the influence of IL-1β on NPCs [10]. However, this low IL-1 receptor expression was surprising in light of previous work showing high *IL1R1* (but not *IL1R2*) expression in rat ventral mesencephalon neural progenitors [10]. It is possible that there is lower *IL1R1* expression in the cortex than in other parts of the fetal brain; or perhaps *IL1R1* expression is upregulated at a later stage of prenatal neurodevelopment in humans than in rats, demonstrating the significance of leveraging human systems in the study of human neurodevelopment. It is also possible that that, by the 24-hour time point, any effect induced by IL-1β may have been lost – i.e., the effects of IL-1β may be very rapid and transient.

Nevertheless, pathway analyses did reveal significant enrichment of gene sets in response to IL-1β treatment, suggesting that there were transcriptional effects in both schizophrenia and control NPCs in response to IL-1β, but our sample size only allowed their identification at the gene set level (at which the multiple-testing burden is smaller). The GSEA analyses revealed substantially different gene set enrichment profiles for the IL-1β signatures in schizophrenia and control NPCs. Among the significantly enriched gene sets in control lines treated with IL-1β were (as with IFNγ) genes regulating the immune response, the presynapse and the post-synaptic density – all upregulated.

However, none of these gene sets were among those most significantly enriched in IL-1β-treated schizophrenia lines; instead, almost all of these were sets of genes involved in central nervous system development and neuronal morphogenesis – all downregulated. The genes that responded differently to IL-1β in schizophrenia vs control NPCs were enriched in gene sets involved in cell division, antigen presentation and, once again, synaptic transmission. The most interesting finding to emerge from all these analyses was indeed that genes involved in synaptic transmission respond differentially to IFNγ and IL-1β exposure in schizophrenia NPCs compared to control NPCs

The current study was primarily limited by the relatively small sample size, which would warrant future replication studies. It was also surprising to find that donor age had some influence over the variance in gene expression in the sample, as one would also expect a negation of age-related epigenetic effects [79]. It is likely that this is due to noise (again as a result of the small sample size), which emphasises the importance of replicating these findings. We also administered only a single, acute dose of cytokine treatment to our NPC cultures: in future studies, it would be interesting to examine the effect of chronic treatment. For IL-1β effects, future studies looking at different developmental time points and cell types could elucidate this cytokine’s role in neurodevelopmental processes associated with schizophrenia.

In summary, having conducted the first transient cytokine exposure study using hiPSC-derived neural progenitor cells from patients with schizophrenia, we have found that immune activation induced by IL-1β and IFNγ elicits transcriptional changes that may alter the course of subsequent neurodevelopment. There were two particularly significant take-home messages from this study, as follows. First, there does appear to be a significant transcriptional response to IFNγ treatment in NPCs, with differential expression implicating mitochondrial complex genes, which are underexpressed in response to treatment in schizophrenia lines. Second, our findings highlight pre- and post-synaptic genes as differentially expressed in response to IFNγ, and differentially regulated in response to treatment in schizophrenia NPCs. In other words, schizophrenia NPCs do not upregulate synaptic genes in response to a cytokine challenge as much as control NPCs do. This is also consistent with previous literature, including large-scale transcriptome-wide association studies [56] and the recent Psychiatric Genetics Consortium study showing synaptic genes to be the most enriched for schizophrenia risk [81]. This is particularly interesting as NPCs do not have synapses. It could be that these early changes impact synaptic development after these cells differentiate into neurons. Indeed, our previous research shows that IFNγ induces molecular and cellular changes in NPCs that persist even when these cells differentiate into neurons [2]. Our findings exemplify differences in how the brains of people with schizophrenia may have responded to infection or inflammation during prenatal development and suggest immune insults early in life can alter neurotransmission. Finally, we identify new gene targets for future research on the influence of maternal immune activation on schizophrenia susceptibility and resilience.

### Author contributions

A.B. wrote the manuscript, conducted the cell culture and experimental treatments, and worked with H.I. on quality control of the genetic data and statistical analyses. H.I. worked on the analyses as well as imputation and quality control of the genetic data and edited the manuscript. P.R., L.D.P and R.N. assisted with the cell culture and experimental treatments. R.R.D conducted the *GeneOverlap* analyses and edited the manuscript. P.J.M.D., C.S., R.N., and D.P.S. set up the experimental pipelines, conducted cell culture and cellular analysis of iPSC and neural progenitor cells, G.M., C.I., R.N. and J.P. were involved in obtaining ethical approval, sample collection and reprogramming of keratinocyte samples. S.B. was responsible for recruitment and clinical evaluation of patients whose hair samples were used in this study. A.V., T.P, E.B., J.P. and D.P.S. edited and provided guidance on the manuscript and interpretation of the results. D.P.S is the principal investigator, corresponding author and PhD supervisor to A.B., oversaw the project and developed the study with A.B., provided essential guidance throughout, as well as writing and editing the manuscript and figures. All of the co-authors provided intellectual input to the study.

## Acknowledgements

We would like to thank all of the participants who took part in this research, as well as the clinical staff who facilitated their involvement. The work (at King’s College, London) was also supported by the Medical Research Council (MRC) Centre grant (MR/N026063/1). A.B. was supported by a Medical Research Council doctoral studentship [MR/N013867/1] for this work. H.I. received funding from the European Union’s Horizon 2020 research and innovation programme under the Marie Skłodowska-Curie grant agreement no. 747429. The study was supported by grants from StemBANCC: the Innovative Medicines Initiative joint undertaking under grant 115439-2, whose resources are composed of financial contribution from the European Union [FP7/2007-2013] and EFPIA companies’ in-kind contribution (JP, DPS); MATRICS: the European Union’s Seventh Framework Programme (FP7-HEALTH-603016) (DPS, JP). In addition, funds from the Wellcome Trust ISSF Grant (No. 097819) and the King’s Health Partners Research and Development Challenge Fund, a fund administered on behalf of King’s Health Partners by Guy’s and St Thomas’ Charity awarded to DPS; the Brain and Behavior Foundation (formally National Alliance for Research on Schizophrenia and Depression (NARSAD); Grant No. 25957), awarded to DPS. E.B. thanks the following funders: the National Institute for Health Research (NIHR200756); Mental Health Research UK John Grace QC Scholarship 2018; BMA Margaret Temple Fellowships 2016 and 2006; Medical Research Council (MRC) and Korean Health Industry Development Institute Partnering Award (MC_PC_16014); MRC New Investigator Award (G0901310); MRC Centenary Award (G1100583); MRC project grant G1100583; a National Institute for Health Research UK post-doctoral fellowship (PDA/02/06/016); the Psychiatry Research Trust; the Schizophrenia Research Fund; the Brain and Behaviour Research Foundation’s NARSAD Young Investigator Awards 2005 and 2008; a Wellcome Trust Research Training Fellowship; Wellcome Trust Case Control Consortium awards (085475/B/08/Z and 085475/Z/08/Z); and the NIHR Biomedical Research Centre for Mental Health at the South London and Maudsley NHS Foundation Trust and Institute of Psychiatry King’s College London. SB has been supported by a National Institute for Health Research (NIHR) Clinician Scientist Award (NIHR CS-11-001) and grants from the Medical Research Council (MR/J012149/1) and NIHR Efficacy and Mechanism Evaluation scheme (16/126/53). The views expressed are those of the authors and not necessarily those of the NHS, the NIHR or the Department of Health. In addition, this paper represents independent research part funded by the National Institute for Health Research (NIHR) Mental Health Biomedical Research Centre (BRC) at South London and Maudsley NHS Foundation Trust and King’s College London. The views expressed are those of the author(s) and not necessarily those of the NHS, the NIHR or the Department of Health and Social Care.

The authors thank the Wohl Cellular Imaging Centre (WCIC) at the IoPPN, King’s College, London, for help with microscopy.

## Data and code availability

RNA sequencing data reported in this paper is available at https://www. ncbi.nlm.nih.gov/subs/sra/SUB10767669. Previously unpublished code used for statistical analyses reported in this paper is available on GitHub: https://github.com/abhat92/DGE_response_to_cytokines_in_SZ_hiPSC-NPCs.

## References

1. Estes ML, McAllister AK: Maternal immune activation: Implications for neuropsychiatric disorders. Science 2016, 353(6301):772–777.

2. Warre-Cornish K, Perfect L, Nagy R, Duarte RRR, Reid MJ, Raval P, Mueller A, Evans AL, Couch A, Ghevaert C et al: Interferon-gamma signaling in human iPSC-derived neurons recapitulates neurodevelopmental disorder phenotypes. Sci Adv 2020, 6(34):eaay9506.

3. Byrne M, Agerbo E, Bennedsen B, Eaton WW, Mortensen PB: Obstetric conditions and risk of first admission with schizophrenia: a Danish national register based study. Schizophr Res 2007, 97(1-3):51–59.

4. Kepinska AP, Iyegbe CO, Vernon AC, Yolken R, Murray RM, Pollak TA: Schizophrenia and Influenza at the Centenary of the 1918-1919 Spanish Influenza Pandemic: Mechanisms of Psychosis Risk. Front Psychiatry 2020, 11:72.

5. Meyer U: Neurodevelopmental Resilience and Susceptibility to Maternal Immune Activation. Trends Neurosci 2019, 42(11):793-806.

6. Garay PA, Hsiao EY, Patterson PH, McAllister AK: Maternal immune activation causes age- and region-specific changes in brain cytokines in offspring throughout development. Brain Behav Immun 2013, 31:54–68.

7. Gilmore JH, Fredrik Jarskog L, Vadlamudi S, Lauder JM: Prenatal infection and risk for schizophrenia: IL-1beta, IL-6, and TNFalpha inhibit cortical neuron dendrite development. Neuropsychopharmacology 2004, 29(7):1221–1229.

8. Gilmore JH, Jarskog LF, Vadlamudi S: Maternal poly I:C exposure during pregnancy regulates TNF alpha, BDNF, and NGF expression in neonatal brain and the maternal-fetal unit of the rat. J Neuroimmunol 2005, 159(1-2):106–112.

9. Lesh TA, Careaga M, Rose DR, McAllister AK, Van de Water J, Carter CS, Ashwood P: Cytokine alterations in first-episode schizophrenia and bipolar disorder: relationships to brain structure and symptoms. J Neuroinflammation 2018, 15(1):165.

10. Crampton SJ, Collins LM, Toulouse A, Nolan YM, O’Keeffe GW: Exposure of foetal neural progenitor cells to IL-1beta impairs their proliferation and alters their differentiation - a role for maternal inflammation? J Neurochem 2012, 120(6):964–973.

11. Goldsmith DR, Rapaport MH, Miller BJ: A meta-analysis of blood cytokine network alterations in psychiatric patients: comparisons between schizophrenia, bipolar disorder and depression. Mol Psychiatry 2016, 21(12):1696–1709.

12. Allswede DM, Yolken RH, Buka SL, Cannon TD: Cytokine concentrations throughout pregnancy and risk for psychosis in adult offspring: a longitudinal case-control study. Lancet Psychiatry 2020, 7(3):254–261.

13. Arrode-Bruses G, Bruses JL: Maternal immune activation by poly I:C induces expression of cytokines IL-1beta and IL-13, chemokine MCP-1 and colony stimulating factor VEGF in fetal mouse brain. J Neuroinflammation 2012, 9:83.

14. Mueller FS, Scarborough J, Schalbetter SM, Richetto J, Kim E, Couch A, Yee Y, Lerch JP, Vernon AC, Weber-Stadlbauer U et al: Behavioral, neuroanatomical, and molecular correlates of resilience and susceptibility to maternal immune activation. Mol Psychiatry 2021, 26(2):396–410.

15. Carlezon WA, Jr., Kim W, Missig G, Finger BC, Landino SM, Alexander AJ, Mokler EL, Robbins JO, Li Y, Bolshakov VY et al: Maternal and early postnatal immune activation produce sex-specific effects on autism-like behaviors and neuroimmune function in mice. Sci Rep 2019, 9(1):16928.

16. Estes ML, Prendergast K, MacMahon JA, Cameron S, Aboubechara JP, Farrelly K, Sell GL, Haapanen L, Schauer JD, Horta A, et al: Baseline immunoreactivity before pregnancy and poly(I:C) dose combine to dictate susceptibility and resilience of offspring to maternal immune activation. Brain Behav Immun 2020, 88:619–630.

17. Brown AS: Prenatal infection as a risk factor for schizophrenia. Schizophr Bull 2006, 32(2):200-202.

18. Brown AS, Begg MD, Gravenstein S, Schaefer CA, Wyatt RJ, Bresnahan M, Babulas VP, Susser ES: Serologic evidence of prenatal influenza in the etiology of schizophrenia. Arch Gen Psychiatry 2004, 61(8):774–780.

19. Feigenson KA, Kusnecov AW, Silverstein SM: Inflammation and the two-hit hypothesis of schizophrenia. Neurosci Biobehav Rev 2014, 38:72–93.

20. van Os J, Rutten BP, Poulton R: Gene-environment interactions in schizophrenia: review of epidemiological findings and future directions. Schizophr Bull 2008, 34(6):1066–1082.

21. Bayer TA, Falkai P, Maier W: Genetic and non-genetic vulnerability factors in schizophrenia: the basis of the "two hit hypothesis". J Psychiatr Res 1999, 33(6):543–548.

22. Leenaars CHC, Kouwenaar C, Stafleu FR, Bleich A, Ritskes-Hoitinga M, De Vries RBM, Meijboom FLB: Animal to human translation: a systematic scoping review of reported concordance rates. J Transl Med 2019, 17(1):223.

23. Pound P, Ritskes-Hoitinga M: Is it possible to overcome issues of external validity in preclinical animal research? Why most animal models are bound to fail. J Transl Med 2018, 16(1):304.

24. Masopust D, Sivula CP, Jameson SC: Of Mice, Dirty Mice, and Men: Using Mice To Understand Human Immunology. J Immunol 2017, 199(2):383–388.

25. Canetta S, Kellendonk C: Can we use mice to study schizophrenia? Philos Trans R Soc Lond B Biol Sci 2018, 373(1742).

26. Takahashi K, Yamanaka S: Induction of pluripotent stem cells from mouse embryonic and adult fibroblast cultures by defined factors. Cell 2006, 126(4):663–676.

27. Aasen T, Izpisua Belmonte JC: Isolation and cultivation of human keratinocytes from skin or plucked hair for the generation of induced pluripotent stem cells. Nat Protoc 2010, 5(2):371–382.

28. Petit I, Kesner NS, Karry R, Robicsek O, Aberdam E, Muller FJ, Aberdam D, Ben-Shachar D: Induced pluripotent stem cells from hair follicles as a cellular model for neurodevelopmental disorders. Stem Cell Res 2012, 8(1):134–140.

29. Brennand K, Savas JN, Kim Y, Tran N, Simone A, Hashimoto-Torii K, Beaumont KG, Kim HJ, Topol A, Ladran I et al: Phenotypic differences in hiPSC NPCs derived from patients with schizophrenia. Mol Psychiatry 2015, 20(3):361–368.

30. Brennand KJ, Gage FH: Modeling psychiatric disorders through reprogramming. Dis Model Mech 2012, 5(1):26–32.

31. Martinez-Cerdeno V, Noctor SC: Neural Progenitor Cell Terminology. Front Neuroanat 2018, 12:104.

32. Shum C, Dutan L, Annuario E, Warre-Cornish K, Taylor SE, Taylor RD, Andreae LC, Buckley NJ, Price J, Bhattacharyya S et al: Delta(9)-tetrahydrocannabinol and 2-AG decreases neurite outgrowth and differentially affects ERK1/2 and Akt signaling in hiPSC-derived cortical neurons. Mol Cell Neurosci 2020, 103:103463.

33. Adhya D, Swarup V, Nagy R, Dutan L, Shum C, Valencia-Alarcon EP, Jozwik KM, Mendez MA, Horder J, Loth E et al: Atypical Neurogenesis in Induced Pluripotent Stem Cells From Autistic Individuals. Biol Psychiatry 2020.

34. Kathuria A, Nowosiad P, Jagasia R, Aigner S, Taylor RD, Andreae LC, Gatford NJF, Lucchesi W, Srivastava DP, Price J: Stem cell-derived neurons from autistic individuals with SHANK3 mutation show morphogenetic abnormalities during early development. Mol Psychiatry 2018, 23(3):735–746.

35. Hoffman GE, Hartley BJ, Flaherty E, Ladran I, Gochman P, Ruderfer DM, Stahl EA, Rapoport J, Sklar P, Brennand KJ: Transcriptional signatures of schizophrenia in hiPSC-derived NPCs and neurons are concordant with post-mortem adult brains. Nat Commun 2017, 8(1):2225.

36. Broadbelt K, Byne W, Jones LB: Evidence for a decrease in basilar dendrites of pyramidal cells in schizophrenic medial prefrontal cortex. Schizophr Res 2002, 58(1):75–81.

37. Ranlund S, Adams RA, Diez A, Constante M, Dutt A, Hall MH, Maestro Carbayo A, McDonald C, Petrella S, Schulze K et al: Impaired prefrontal synaptic gain in people with psychosis and their relatives during the mismatch negativity. Hum Brain Mapp 2016, 37(1):351–365.

38. Baines KJ, Hillier DM, Haddad FL, Rajakumar N, Schmid S, Renaud SJ: Maternal Immune Activation Alters Fetal Brain Development and Enhances Proliferation of Neural Precursor Cells in Rats. Front Immunol 2020, 11:1145.

39. World Health O: International statistical classification of diseases and related health problems : tenth revision. Geneva: World Health Organization; 2004.

40. McGuffin P, Farmer A, Harvey I: A polydiagnostic application of operational criteria in studies of psychotic illness. Development and reliability of the OPCRIT system. Arch Gen Psychiatry 1991, 48(8):764–770.

41. Cocks G, Curran S, Gami P, Uwanogho D, Jeffries AR, Kathuria A, Lucchesi W, Wood V, Dixon R, Ogilvie C et al: The utility of patient specific induced pluripotent stem cells for the modelling of Autistic Spectrum Disorders. Psychopharmacology (Berl) 2014, 231(6):1079–1088.

42. Sheridan SD, Surampudi V, Rao RR: Analysis of embryoid bodies derived from human induced pluripotent stem cells as a means to assess pluripotency. Stem Cells Int 2012, 2012:738910.

43. Chambers I, Tomlinson SR: The transcriptional foundation of pluripotency. Development 2009, 136(14):2311–2322.

44. International Stem Cell Initiative, Adewumi O, Aflatoonian B, Ahrlund-Richter L, …, Weaver TA, Young LA, Zhang W: Characterization of human embryonic stem cell lines by the International Stem Cell Initiative. Nat Biotechnol 2007, 25(7):803–816.

45. Boulting GL, Kiskinis E, Croft GF, Amoroso MW, Oakley DH, Wainger BJ, Williams DJ, Kahler DJ, Yamaki M, Davidow L et al: A functionally characterized test set of human induced pluripotent stem cells. Nat Biotechnol 2011, 29(3):279–286.

46. Muller FJ, Schuldt BM, Williams R, Mason D, Altun G, Papapetrou EP, Danner S, Goldmann JE, Herbst A, Schmidt NO et al: A bioinformatic assay for pluripotency in human cells. Nat Methods 2011, 8(4):315–317.

47. Baruzzo G, Hayer KE, Kim EJ, Di Camillo B, FitzGerald GA, Grant GR: Simulation-based comprehensive benchmarking of RNA-seq aligners. Nat Methods 2017, 14(2):135–139.

48. Liao Y, Smyth GK, Shi W: featureCounts: an efficient general purpose program for assigning sequence reads to genomic features. Bioinformatics 2014, 30(7):923–930.

49. Law CW, Chen Y, Shi W, Smyth GK: voom: Precision weights unlock linear model analysis tools for RNA-seq read counts. Genome Biol 2014, 15(2):R29.

50. Hoffman GE, Schadt EE: variancePartition: interpreting drivers of variation in complex gene expression studies. BMC Bioinformatics 2016, 17(1):483.

51. Hoffman GE, Roussos P: dream: Powerful differential expression analysis for repeated measures designs. Bioinformatics 2020.

52. Schrode N, Ho SM, Yamamuro K, Dobbyn A, Huckins L, Matos MR, Cheng E, Deans PJM, Flaherty E, Barretto N et al: Synergistic effects of common schizophrenia risk variants. Nat Genet 2019, 51(10):1475–1485.

53. Mertens J, Paquola ACM, Ku M, Hatch E, Bohnke L, Ladjevardi S, McGrath S, Campbell B, Lee H, Herdy JR et al: Directly Reprogrammed Human Neurons Retain Aging-Associated Transcriptomic Signatures and Reveal Age-Related Nucleocytoplasmic Defects. Cell Stem Cell 2015, 17(6):705–718.

54. Pocklington AJ, Rees E, Walters JT, Han J, Kavanagh DH, Chambert KD, Holmans P, Moran JL, McCarroll SA, Kirov G et al: Novel Findings from CNVs Implicate Inhibitory and Excitatory Signaling Complexes in Schizophrenia. Neuron 2015, 86(5):1203–1214.

55. Pardiñas AF, Holmans P, Pocklington AJ, Escott-Price V, Ripke S, Carrera N, Legge SE, Bishop S, Cameron D, Hamshere ML et al: Common schizophrenia alleles are enriched in mutation-intolerant genes and in regions under strong background selection. Nat Genet 2018, 50(3):381–389.

56. Hall LS, Medway CW, Pain O, Pardinas AF, Rees EG, Escott-Price V, Pocklington A, Bray NJ, Holmans PA, Walters JTR et al: A transcriptome-wide association study implicates specific pre- and post-synaptic abnormalities in schizophrenia. Hum Mol Genet 2020, 29(1):159–167.

57. Bhat A, Irizar H, Thygesen JH, Kuchenbaecker K, Pain O, Adams RA, Zartaloudi E, Harju-Seppanen J, Austin-Zimmerman I, Wang B et al: Transcriptome-wide association study reveals two genes that influence mismatch negativity. Cell Rep 2021, 34(11):108868.

58. Korotkevich G, Sukhov V, Sergushichev A: Fast gene set enrichment analysis. In: bioRxiv. 2019.

59. Gandal MJ, Zhang P, Hadjimichael E, Walker RL, Chen C, Liu S, Won H, van Bakel H, Varghese M, Wang Y et al: Transcriptome-wide isoform-level dysregulation in ASD, schizophrenia, and bipolar disorder. *Science (New York*, NY) 2018, 362(6420).

60. Pardiñas AF, Holmans P, Pocklington AJ, Escott-Price V, Ripke S, Carrera N, Legge SE, Bishop S, Cameron D, Hamshere ML et al: Common schizophrenia alleles are enriched in mutation-intolerant genes and in regions under strong background selection. Nature Genetics 2018, 50(3):381–389.

61. de Leeuw CA, Mooij JM, Heskes T, Posthuma D: MAGMA: Generalized Gene-Set Analysis of GWAS Data. PLOS Comp Bio 2015, 11(4):e1004219.

62. Majoros A, Platanitis E, Kernbauer-Holzl E, Rosebrock F, Muller M, Decker T: Canonical and Non-Canonical Aspects of JAK-STAT Signaling: Lessons from Interferons for Cytokine Responses. Front Immunol 2017, 8:29.

63. Honkala AT, Tailor D, Malhotra SV: Guanylate-Binding Protein 1: An Emerging Target in Inflammation and Cancer. Front Immunol 2019, 10:3139.

64. Gandal MJ, Zhang P, Hadjimichael E, Walker RL, Chen C, Liu S, Won H, van Bakel H, Varghese M, Wang Y et al: Transcriptome-wide isoform-level dysregulation in ASD, schizophrenia, and bipolar disorder. Science 2018, 362(6420).

65. Shen L: GeneOverlap: Test and visualize gene overlaps. R package version 1.26.0. . 2020.

66. Pain O, Pocklington AJ, Holmans PA, Bray NJ, O’Brien HE, Hall LS, Pardinas AF, O’Donovan MC, Owen MJ, Anney R: Novel Insight Into the Etiology of Autism Spectrum Disorder Gained by Integrating Expression Data With Genome-wide Association Statistics. Biol Psychiatry 2019, 86(4):265–273.

67. Greenhalgh AD, David S, Bennett FC: Immune cell regulation of glia during CNS injury and disease. Nat Rev Neurosci 2020, 21(3):139–152.

68. Park GH, Noh H, Shao Z, Ni P, Qin Y, Liu D, Beaudreault CP, Park JS, Abani CP, Park JM et al: Activated microglia cause metabolic disruptions in developmental cortical interneurons that persist in interneurons from individuals with schizophrenia. Nat Neurosci 2020, 23(11):1352–1364.

69. Tang BM, Shojaei M, Parnell GP, Huang S, Nalos M, Teoh S, O’Connor K, Schibeci S, Phu AL, Kumar A et al: A novel immune biomarker IFI27 discriminates between influenza and bacteria in patients with suspected respiratory infection. Eur Respir J 2017, 49(6).

70. Francisco LM, Sage PT, Sharpe AH: The PD-1 pathway in tolerance and autoimmunity. Immunol Rev 2010, 236:219–242.

71. Olaya JC, Heusner CL, Matsumoto M, Sinclair D, Kondo MA, Karl T, Shannon Weickert C: Overexpression of Neuregulin 1 Type III Confers Hippocampal mRNA Alterations and Schizophrenia-Like Behaviors in Mice. Schizophr Bull 2018, 44(4):865–875.

72. Malhotra D, McCarthy S, Michaelson JJ, Vacic V, Burdick KE, Yoon S, Cichon S, Corvin A, Gary S, Gershon ES et al: High frequencies of de novo CNVs in bipolar disorder and schizophrenia. Neuron 2011, 72(6):951–963.

73. Rajasekaran A, Venkatasubramanian G, Berk M, Debnath M: Mitochondrial dysfunction in schizophrenia: pathways, mechanisms and implications. Neurosci Biobehav Rev 2015, 48:10–21.

74. Adams RA, Stephan KE, Brown HR, Frith CD, Friston KJ: The computational anatomy of psychosis. Front Psychiatry 2013, 4:47.

75. Friston, Brown HR, Siemerkus J, Stephan KE: The dysconnection hypothesis (2016). Schizophr Res 2016, 176(2-3):83–94.

76. Stephan KE, Friston KJ, Frith CD: Dysconnection in schizophrenia: from abnormal synaptic plasticity to failures of self-monitoring. Schizophr Bull 2009, 35(3):509–527.

77. Stephan KE, Baldeweg T, Friston KJ: Synaptic plasticity and dysconnection in schizophrenia. Biol Psychiatry 2006, 59(10):929–939.

78. Glantz LA, Lewis DA: Reduction of synaptophysin immunoreactivity in the prefrontal cortex of subjects with schizophrenia. Regional and diagnostic specificity. Arch Gen Psychiatry 1997, 54(10):943–952.

79. Steg LC, Shireby GL, Imm J, Davies JP, Franklin A, Flynn R, Namboori SC, Bhinge A, Jeffries AR, Burrage J et al: Novel epigenetic clock for fetal brain development predicts prenatal age for cellular stem cell models and derived neurons. Mol Brain 2021, 14(1):98.

80. Kushima I, Aleksic B, Nakatochi M, Shimamura T, Shiino T, Yoshimi A, Kimura H, Takasaki Y, Wang C, Xing J et al: High-resolution copy number variation analysis of schizophrenia in Japan. Mol Psychiatry 2017, 22(3):430–440.

81. Ripke S, Walters JT, O’Donovan MC: Mapping genomic loci prioritises genes and implicates synaptic biology in schizophrenia. medRxiv 2020:2020.2009.2012.20192922.

82. Shatz, C. J. MHC class I: an unexpected role in neuronal plasticity. Neuron 2009, 64(1), 40

