## Supplemental Information for "Attenuated transcriptional response to pro-inflammatory cytokines in schizophrenia hiPSC-derived neural progenitor cells"

**Supplementary information, tables and figures.**

**Supplementary Tables 1 & 3**

**Supplementary Figures 1 - 9**

**Supplementary Tables**

**Supplementary Table 1.** Top 20 genes most significantly differentially expressed in IFNγ-treated control cells compared to untreated control cells.

| Gene Symbol | Log fold change | Average expression[2] | P value | Adjusted P value | Z score | FDR[1] |
|---|---|---|---|---|---|---|
| IFI27 | 6.067 | 4.280 | 2.16E-10 | 2.97E-06 | 6.350 | 2.97E-06 |
| CD274 | 6.386 | 1.547 | 3.95E-10 | 2.97E-06 | 6.256 | 2.97E-06 |
| IRF1 | 7.022 | 4.265 | 6.44E-10 | 3.24E-06 | 6.179 | 3.23E-06 |
| PSMB10 | 5.522 | 1.528 | 1.15E-09 | 4.32E-06 | 6.087 | 4.33E-06 |
| SP140L | 5.815 | 0.465 | 1.81E-09 | 5.05E-06 | 6.014 | 5.05E-06 |
| STAT2 | 2.247 | 6.869 | 2.01E-09 | 5.05E-06 | 5.997 | 5.05E-06 |
| STAT1 | 5.680 | 7.882 | 2.74E-09 | 5.58E-06 | 5.946 | 5.57E-06 |
| TAP1 | 7.167 | 4.928 | 3.18E-09 | 5.58E-06 | 5.922 | 5.57E-06 |
| GSTK1 | 1.608 | 5.890 | 3.47E-09 | 5.58E-06 | 5.908 | 5.57E-06 |
| PSMB9 | 9.482 | 2.362 | 3.70E-09 | 5.58E-06 | 5.897 | 5.57E-06 |
| ISG15 | 4.980 | 4.931 | 7.22E-09 | 9.89E-06 | 5.786 | 9.88E-06 |
| TNFRSF14 | 4.751 | 1.658 | 8.30E-09 | 1.04E-05 | 5.762 | 1.04E-05 |
| SECTM1 | 5.813 | 0.609 | 9.07E-09 | 1.05E-05 | 5.747 | 1.05E-05 |
| WDFY1 | 1.922 | 7.012 | 1.94E-08 | 2.09E-05 | 5.617 | 2.09E-05 |
| LAP3 | 4.433 | 6.512 | 2.85E-08 | 2.86E-05 | 5.551 | 2.86E-05 |

| | | | | | | |
|---|---|---|---|---|---|---|
| IL18BP | 3.892 | 4.188 | 3.13E-08 | 2.95E-05 | 5.534 | 2.95E-05 |
| GBP1 | 14.622 | 2.693 | 3.51E-08 | 3.11E-05 | 5.514 | 3.11E-05 |
| AL157871.2 | 6.200 | -0.603 | 3.88E-08 | 3.25E-05 | 5.496 | 3.25E-05 |
| XAF1 | 10.252 | -1.583 | 5.23E-08 | 4.15E-05 | 5.443 | 4.15E-05 |

[1] False Discovery Rate

[2] Average expression of the gene in TMM-normalised $\log_2$ CPMs (counts-per-million).

(Please see Supplementary Spreadsheets 3A-G for all genes included in the differential expression comparisons.)

**Supplementary Table 2.** Top 20 genes most significantly differentially expressed in IFNγ-treated schizophrenia cell lines compared to untreated schizophrenia cell lines.

| Gene Symbol | Log fold change | Average expression[2] | P value | Adjusted P value | Z score | FDR[1] |
|---|---|---|---|---|---|---|
| STAT2 | 2.610 | 6.869 | 9.67E-10 | 1.46E-05 | 6.115 | 1.46E-05 |
| IFI27 | 6.333 | 4.280 | 4.00E-09 | 1.74E-05 | 5.884 | 1.74E-05 |
| STAT1 | 5.454 | 7.882 | 4.90E-09 | 1.74E-05 | 5.851 | 1.74E-05 |
| PSMB10 | 5.740 | 1.528 | 5.49E-09 | 1.74E-05 | 5.832 | 1.74E-05 |
| GSTK1 | 1.552 | 5.890 | 5.77E-09 | 1.74E-05 | 5.823 | 1.74E-05 |
| SP140L | 6.125 | 0.465 | 8.51E-09 | 2.14E-05 | 5.758 | 2.14E-05 |
| IRF1 | 7.278 | 4.265 | 1.65E-08 | 3.54E-05 | 5.646 | 3.54E-05 |
| GBP1 | 12.308 | 2.693 | 2.08E-08 | 3.91E-05 | 5.606 | 3.91E-05 |
| ISG15 | 5.142 | 4.931 | 2.92E-08 | 4.88E-05 | 5.546 | 4.88E-05 |
| MT2A | 3.091 | 4.452 | 3.97E-08 | 5.57E-05 | 5.492 | 5.57E-05 |
| TNFRSF14 | 5.052 | 1.658 | 4.07E-08 | 5.57E-05 | 5.488 | 5.57E-05 |
| IFI6 | 4.015 | 6.238 | 4.57E-08 | 5.73E-05 | 5.467 | 5.73E-05 |
| ADAR | 1.182 | 8.246 | 5.86E-08 | 6.02E-05 | 5.423 | 6.02E-05 |
| SECTM1 | 7.449 | 0.609 | 6.01E-08 | 6.02E-05 | 5.419 | 6.02E-05 |
| ERAP2 | 5.936 | 2.491 | 6.23E-08 | 6.02E-05 | 5.412 | 6.02E-05 |
| ITK | 8.431 | -1.891 | 6.66E-08 | 6.02E-05 | 5.400 | 6.02E-05 |

| | | | | | | |
|---|---|---|---|---|---|---|
| MMP25-AS1 | 3.487 | 2.799 | 6.80E-08 | 6.02E-05 | 5.396 | 6.02E-05 |
| XAF1 | 10.277 | -1.583 | 8.11E-08 | 6.79E-05 | 5.365 | 6.79E-05 |
| LAP3 | 4.017 | 6.512 | 9.33E-08 | 7.40E-05 | 5.339 | 7.40E-05 |

[1] False Discovery Rate

[2] Average expression of the gene in TMM-normalised $log_2$ CPMs (counts-per-million).

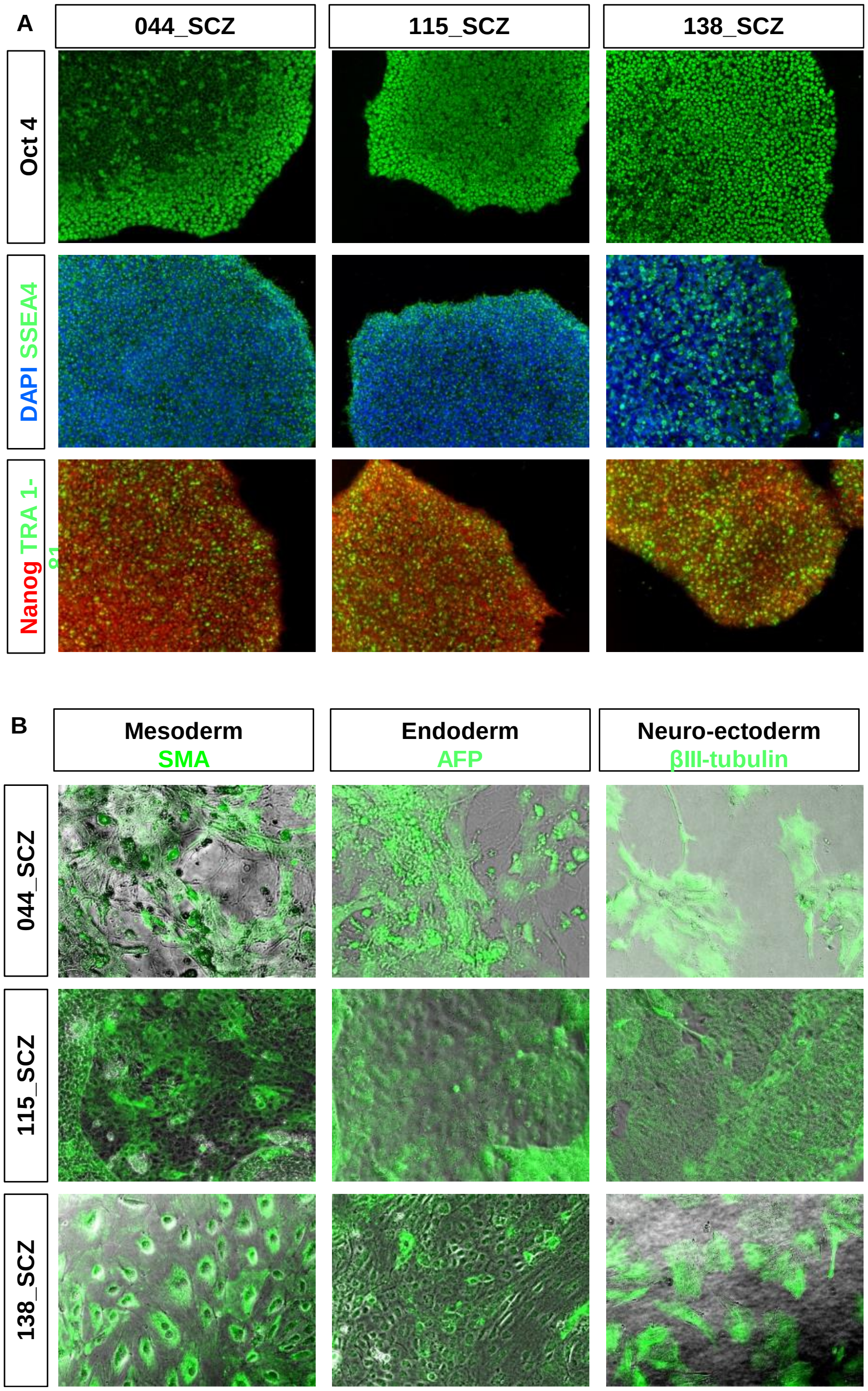


**Supplementary Figure 1.** *Validation of schizophrenia hiPSC lines.* **A.** Shows immunostaining of iPSCs for the pluripotency markers OCT4, SSEA4, NANOG and TRA-1-81. **B.** hiPSCs were further tested for their ability to generate cells from the 3 germ layers, as in Cocks et al (2014). All three of these hiPSC lines showed spontaneous generation of mesoderm, endoderm and neuro-ectoderm cells. Validation of control lines used in this study has been shown in (Adhya et al., 2020; Shum et al., 2020).

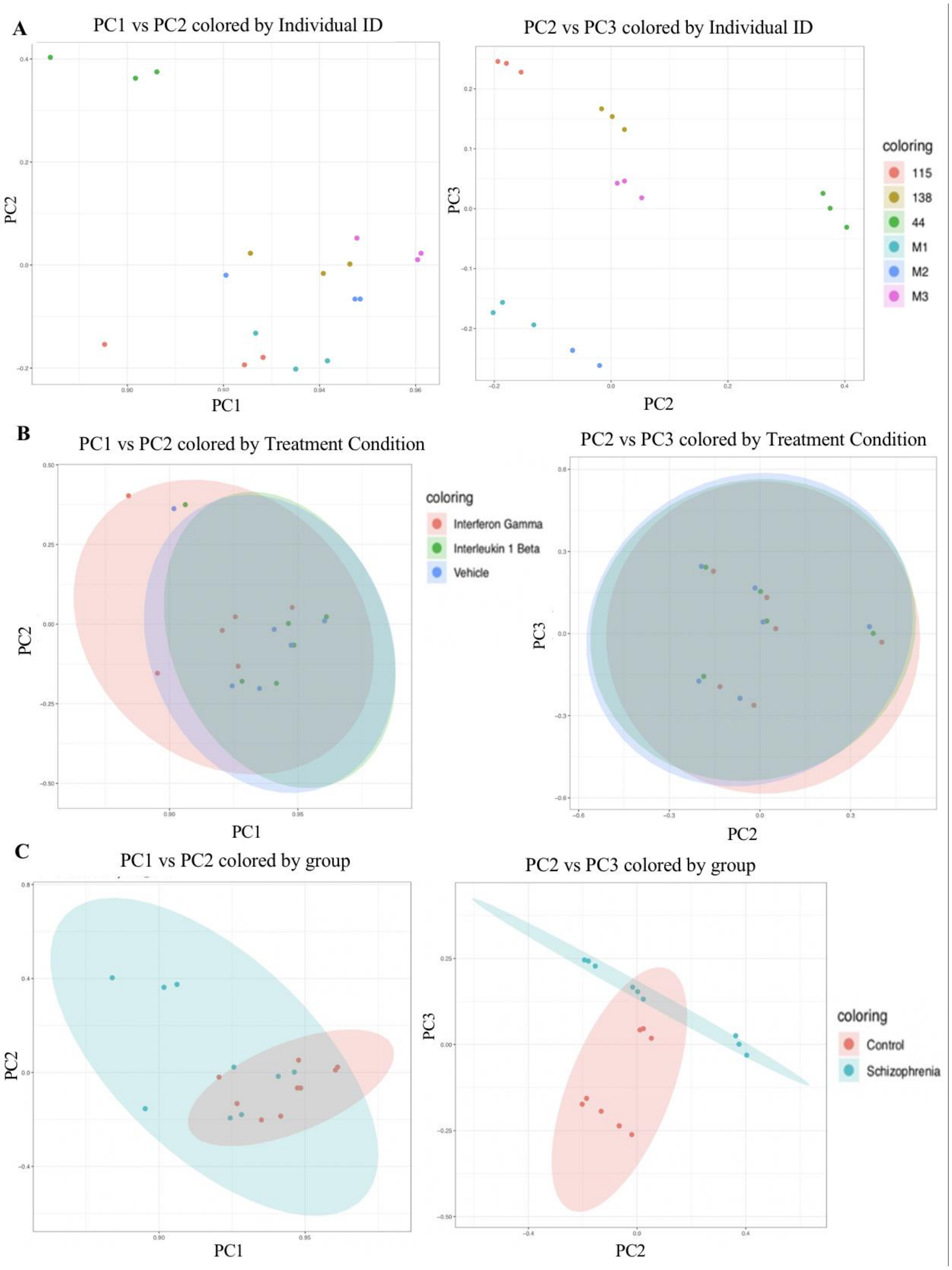


**Supplementary Figure 2.** *Principal component analysis (PCA).* The left panels show the relationship between PC1 and PC2; the right shows PC2 against PC3 (PC1 accounts for 86.55% of the variation in the data). **A.** By individual identifier. **B.** By treatment condition: IFNγ, IL-1β and vehicle (untreated). **C.** By diagnostic group (schizophrenia versus controls).

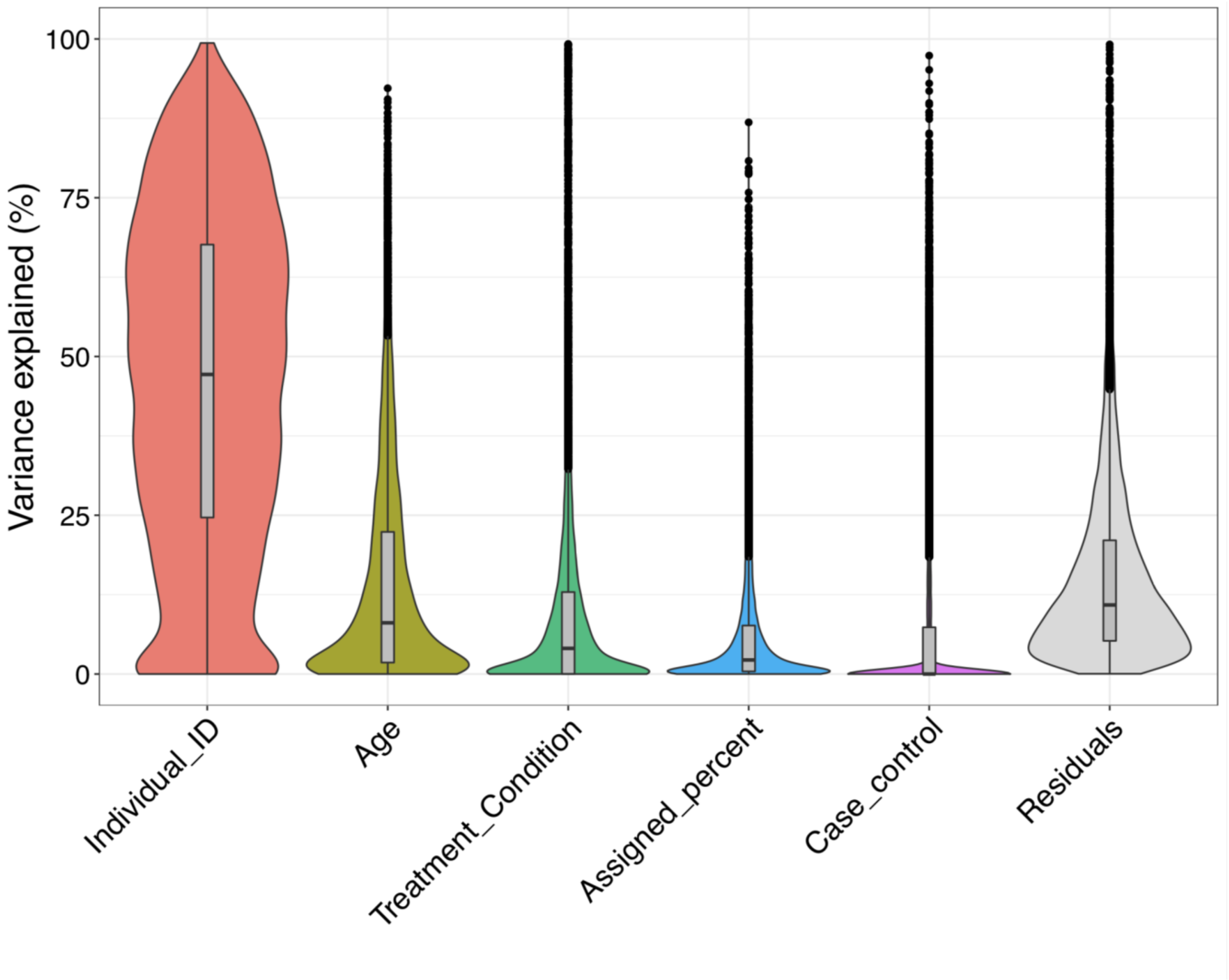


**Supplementary Figure 3.** *Variance partition analysis*. This shows the percentage of variance in the expression of each gene explained by each variable. Due to small sample size, this estimation may be very noisy. This figure mainly serves to illustrate that the primary source of variance in the data is the individual differences between each of the cell lines.

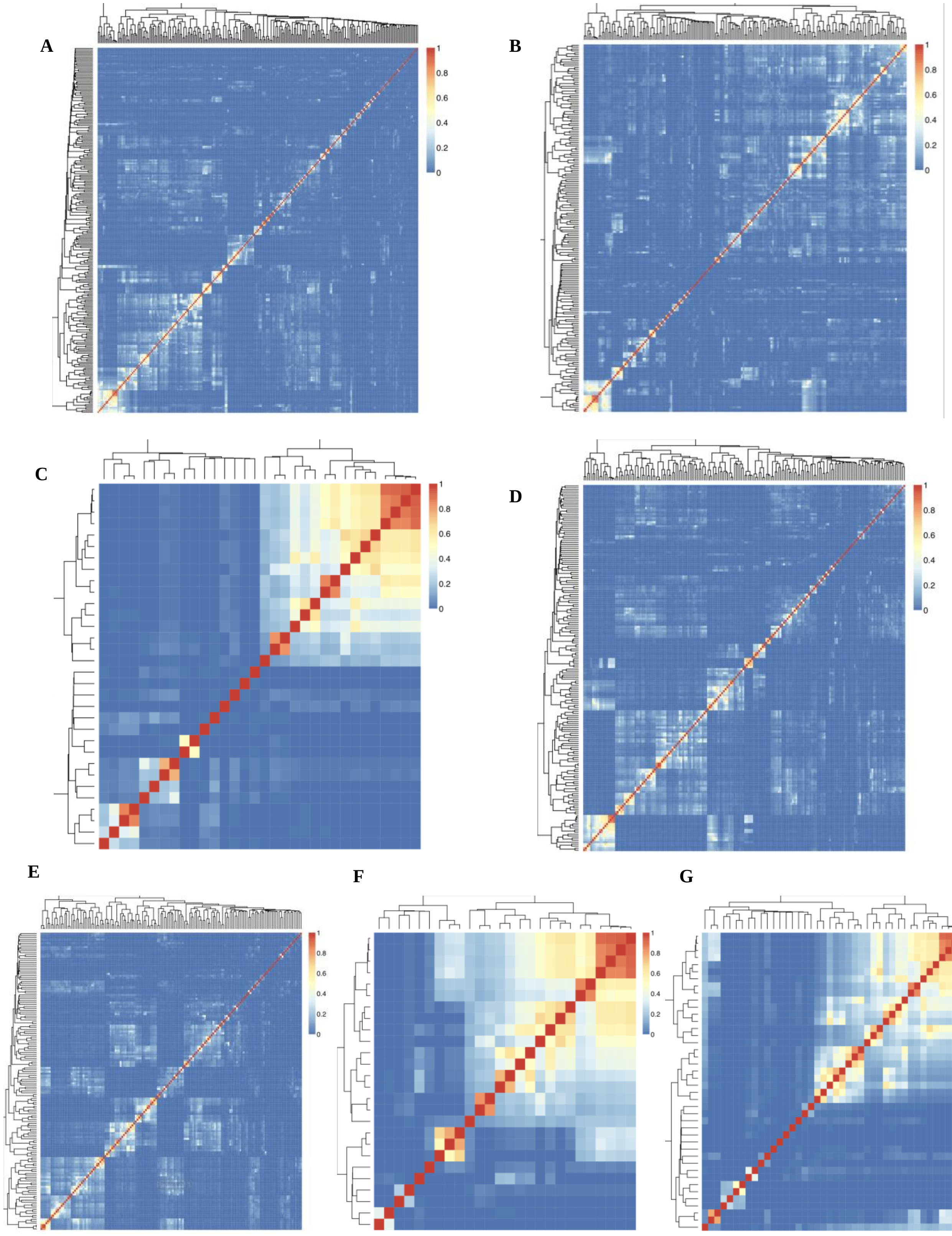


**Supplementary Figure 4.** *Jaccard similarity matrices* for signatures A-G, respectively

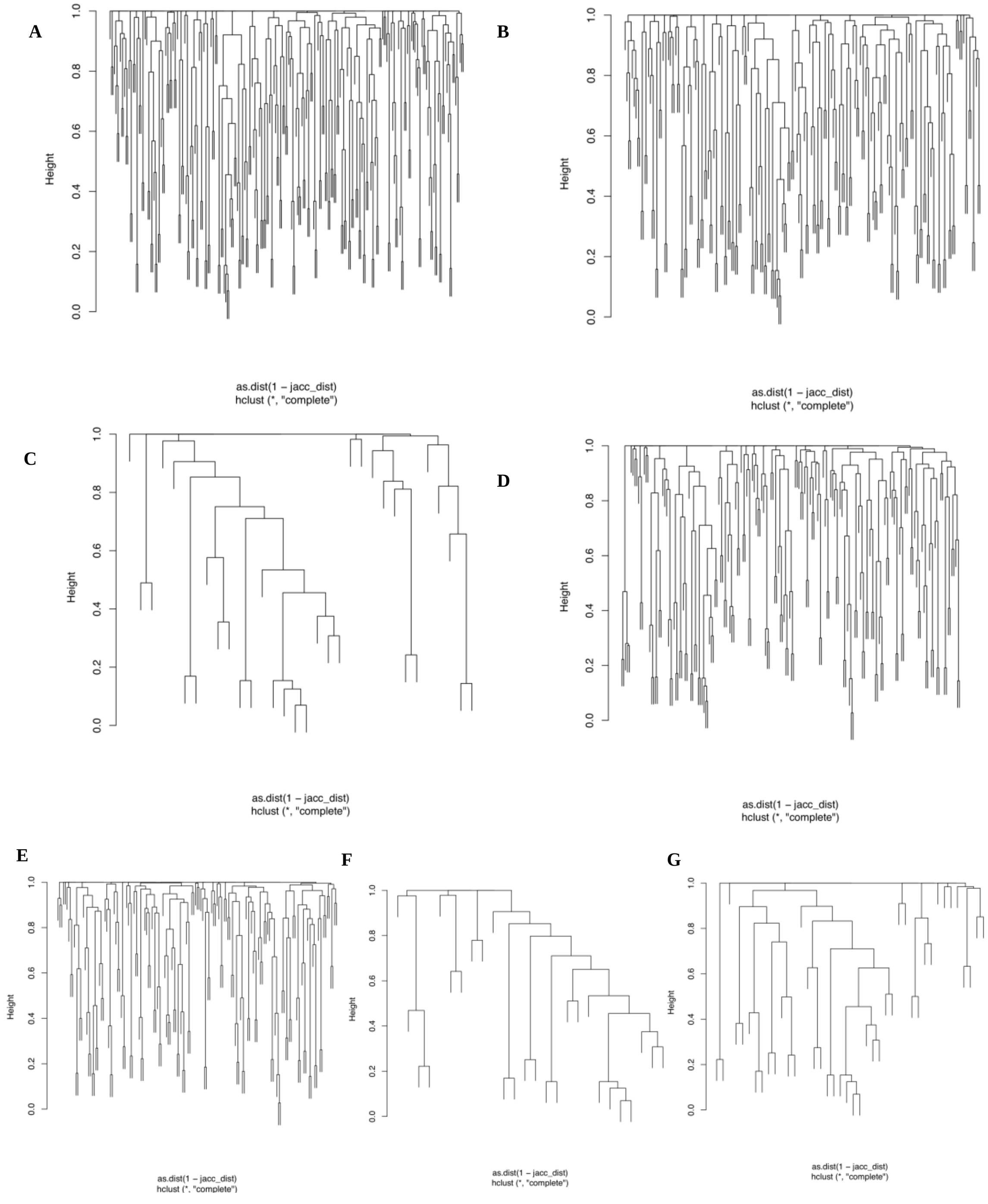


**Supplementary Figure 5.** *Cluster dendograms* for signatures A-G, respectively.

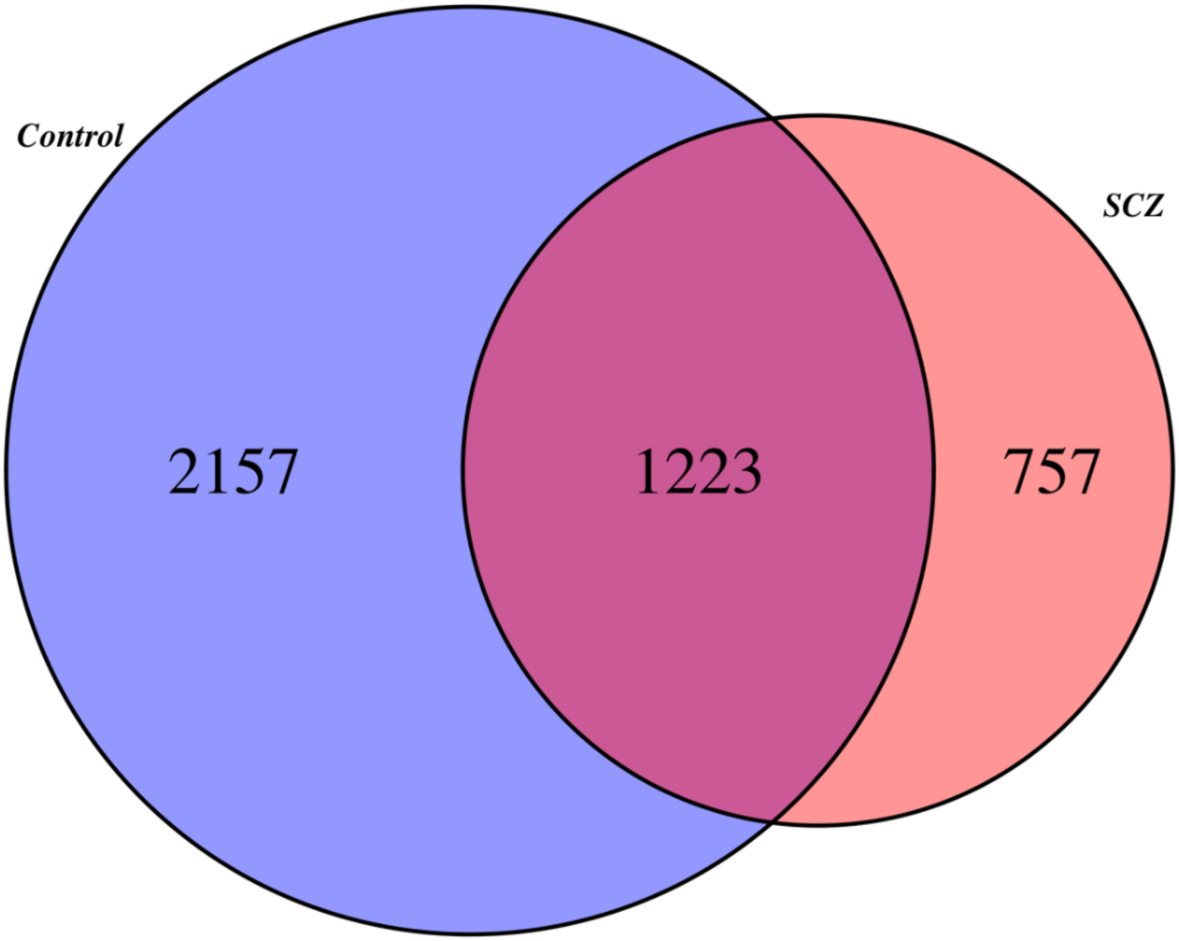


**Supplementary Figure 6.** *Venn diagram that shows the overlap between the genes that are differentially expressed in response to IFNγ in control cells and in schizophrenia cells.* Of the 4137 genes that respond to IFNγ in any of the two groups, only 1223 genes are in common, meaning that there are 2914 genes that appear differentially expressed in response to IFNγ in controls vs not in schizophrenia and vice versa.

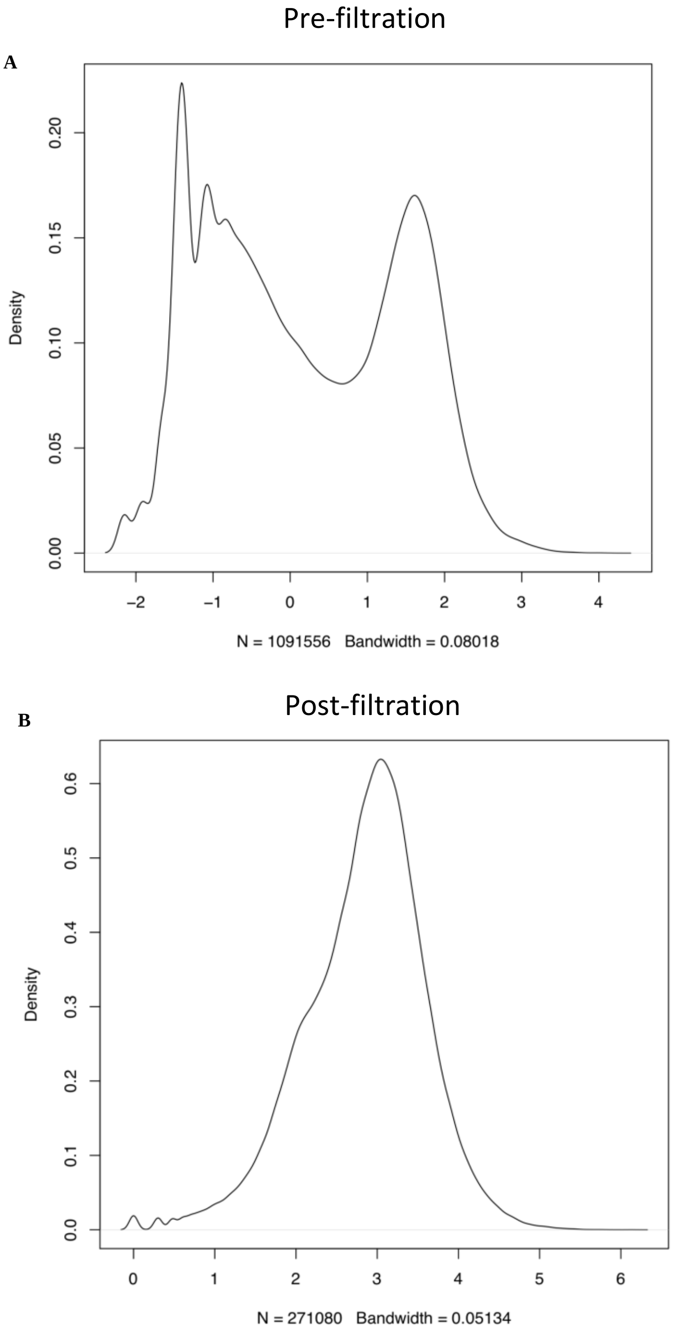


**Supplementary Figure 7.** *Filtration to remove lowly expressed genes in order to minimise technical noise.* Plots **A** and **B** show the distribution of $log_{10}$-transformed counts-per-million (CPMs) pre-filtration and post-filtration, respectively. Filtration threshold was set at $log_{10}CPM = 0.6$ (CPM x $10^{0.6}$) by visual inspection, such that the filtered data showed an approximate Gaussian distribution.

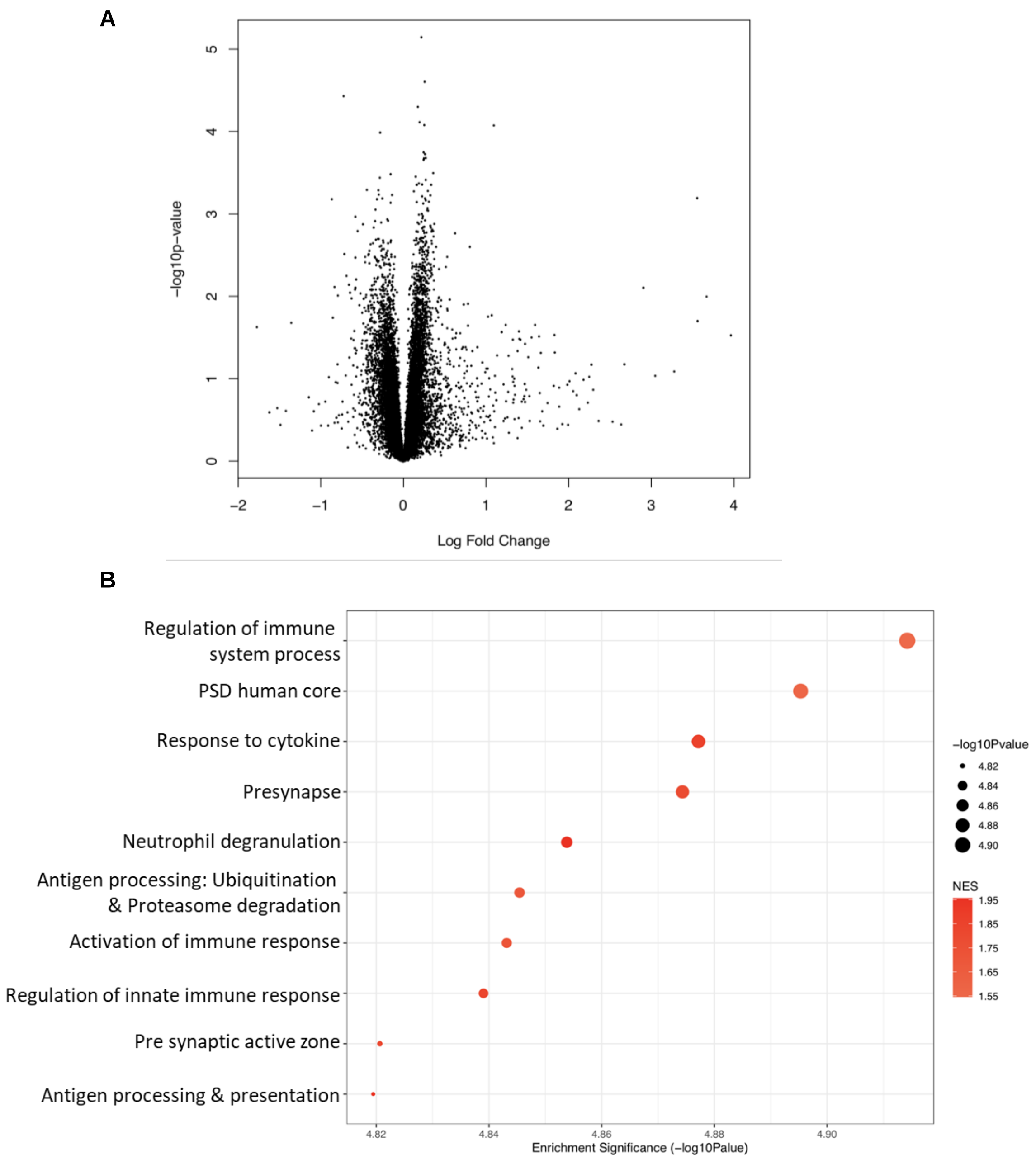


**Supplementary Figure 8. Expression differences between NPCs from IL-1β treated versus untreated control NPCs (signature E) at the gene and pathway level. A.** The volcano plot shows, on the *y*-axis, the statistical significance (-$\log_{10}$ p-value) of differential expression of genes in IL-1β-treated compared to untreated control cells. The *x*-axis is the magnitude of change ($\log_2$ fold change) in expression of those genes after IL-1β treatment. **B.** The top 10 significantly enriched gene set clusters (the gene set with the lowest *p*-value in each cluster is labelled on the *y*-axis). Data-points are sized here according to significance ($-\log_{10}$ p-value), and coloured according to the normalised enrichment score (NES), with darker red indicating greater upregulation.

A

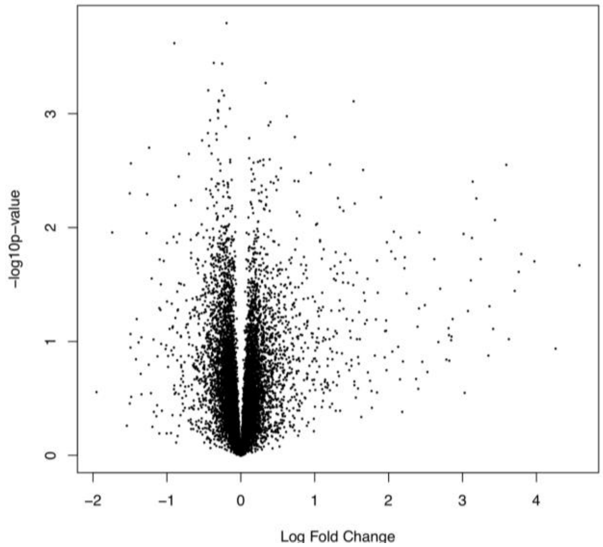


B

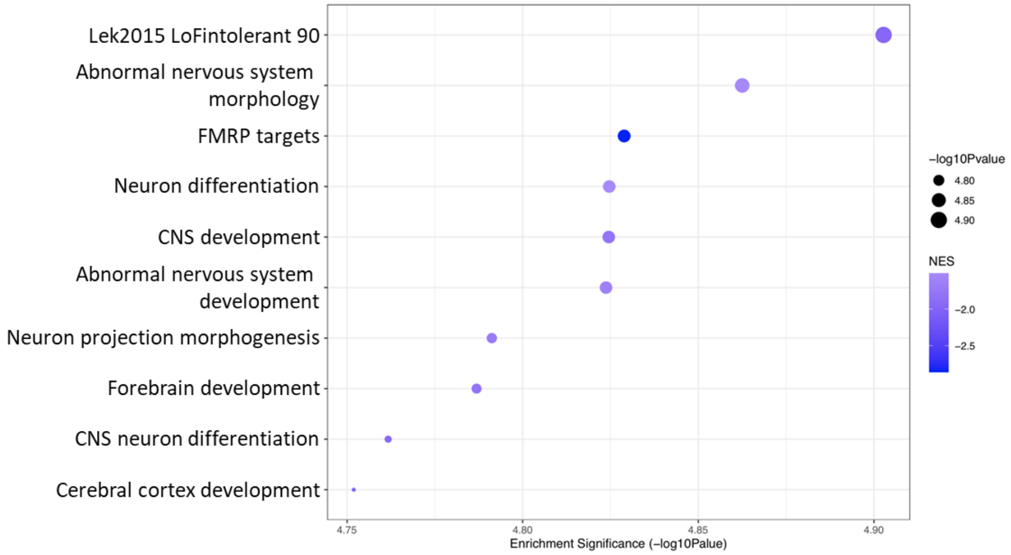


**Supplementary Figure 9. Expression differences between NPCs from IL-1β treated versus untreated schizophrenia NPCs (signature F) at the gene and pathway level. A.** The volcano plot shows, on the *y*-axis, the statistical significance (-$\log_{10}$ p-value) of differential expression of genes in IL-1β-treated compared to untreated cells from patient (SCZ) donors. The *x*-axis is the magnitude of change ($\log_2$ fold change) in expression of those genes after IL-1β treatment. **B.** The top 10 significantly enriched gene set clusters (the gene set with the lowest *p*-value in each cluster is labelled on the *y*-axis). Data-points are sized according to significance ($-\log_{10}$ p-value) and coloured according to normalised enrichment score (NES), with darker blue indicating greater downregulation.

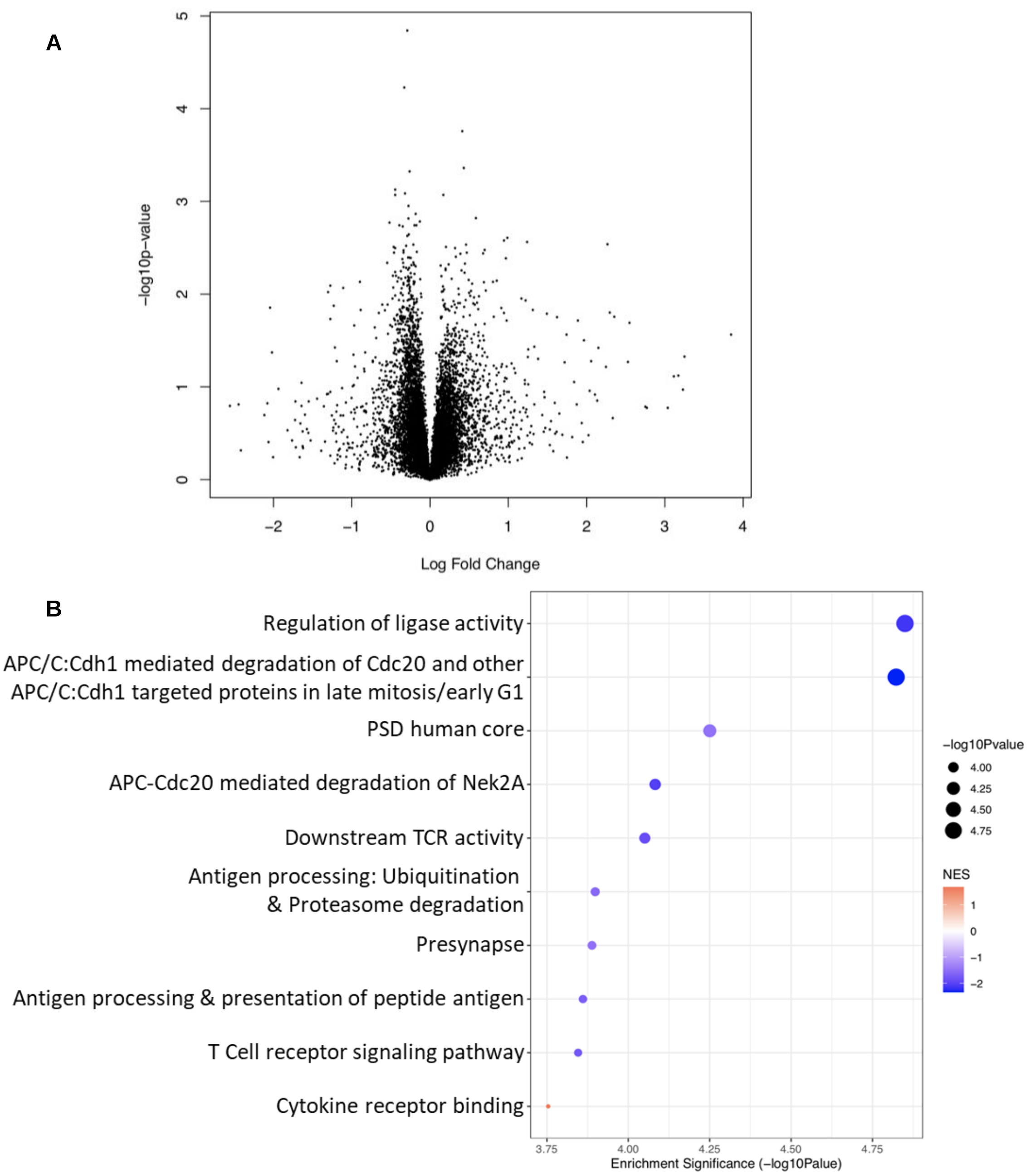


**Supplementary Figure 10. Interaction effect between IL-1β-treatment and diagnostic group on gene expression (Signature G). A.** The volcano plot shows, on the *y*-axis, the statistical significance (-$\log_{10}$ p-value) of differential expression of genes in IL-1β-treated compared to untreated cells from patient donors versus those from control donors (i.e., the interaction effect between IL-1β -treatment and diagnostic group on gene expression). The *x*-axis is the magnitude of change ($\log_2$ fold change) in expression of those genes after IL-1β treatment. **B.** The top ten significantly enriched gene set clusters (the gene set with the lowest *p*-value in each cluster is labelled on the *x*-axis). Data-points are sized according to significance ($-\log_{10}$ p-value) and coloured according to normalised enrichment score (NES), with blue indicating downregulation and red indicating upregulation.
